# Condensin pinches a short negatively supercoiled DNA loop during each round of ATP usage

**DOI:** 10.1101/2022.06.03.494647

**Authors:** Belén Martínez-García, Sílvia Dyson, Joana Segura, Pilar Gutierrez-Escribano, Luís Aragón, Joaquim Roca

## Abstract

Condensin extrudes DNA loops using an ATP-dependent mechanism that remains to be elucidated. Here, we show how condensin activity alters the topology of the interacting DNA. High condensin concentrations restrain DNA positive supercoils. However, in experimental conditions that sustain DNA loop extrusion, condensin restrains negative supercoils. Namely, upon ATP-mediated loading onto DNA, each condensin constrains a DNA linking number difference (ΔLk) of -0.4. This ΔLk increases to -0.8 during ATP binding and resets to -0.4 upon ATP hydrolysis. These ΔLk values reflect the transient formation of a short left-handed loop of DNA, which is not the extruding loop. We conclude that, upon condensin ATPase-head engagement, a segment of DNA is pinched to form a short negatively supercoiled loop, which can be subsequently merged with the extruding loop. Such “pinch and merge” mechanism implies that the DNA is transferred between two dynamic DNA-binding sites while anchored at a third site.

## INTRODUCTION

Structural maintenance of chromosomes (SMC) complexes play key roles in the macroscale architecture and dynamics of chromosomes in all domains of life. In bacteria, SMC-ScpAB and MukBEF promote individualization and segregation of replicated chromosomes (Gruber, 2018; Makela and Sherratt, 2020). In eukaryotes, condensin folds chromatin fibers into rod-shaped chromatids during mitosis, cohesin mediates sister chromatid cohesion and interphase organisation of chromatin, and Smc5/6 is involved in DNA repair (Hirano, 2016; Uhlmann, 2016; Yatskevich et al., 2019). Despite these different roles, all SMC complexes have similar ATPase domains and a common architecture, implying they share a core mechanism of action (Hassler et al., 2018). In this respect, *in vivo, in vitro* and *in silico* studies have converged on the idea that SMC complexes are universal DNA loop extrusion motors (Datta et al., 2020; Davidson and Peters, 2021; Higashi and Uhlmann, 2022; van Ruiten and Rowland, 2018). Supporting this notion, biochemical reconstitution assays have proven that condensin and cohesin can extrude DNA loops at high speed (hundreds of bp/s) by consuming little amounts of ATP (Davidson et al., 2019; Ganji et al., 2018; Kim et al., 2019). How SMC complexes dynamically manipulate DNA fibers to extrude DNA loops remains to be elucidated.

The core SMC complex is a large heterotrimeric protein ring formed by two Smc subunits and a kleisin (Gruber et al., 2003; Haering et al., 2002; Schleiffer et al., 2003). In the budding yeast condensin, these are named Smc2, Smc4 and Brn1, respectively (Figure S1A). Each Smc subunit folds into a 50 nm long antiparallel coiled-coil that forms a globular “hinge” domain at its apex, whereas the amino and carboxy termini form an ABC-type ATPase “head” domain at the other extreme. Smc2 and Smc4 stably dimerize via their hinge domains, while the long and flexible kleisin subunit Brn1 closes the tripartite ring by connecting the two head domains in an asymmetric way. The N-terminal domain of Brn1 binds to the coiled-coil “neck” region immediately adjacent to the head of Smc2, while the C-terminal domain binds to the head tip of Smc4 at a site called the “cap”. The condensin complex is completed by two HEAT repeat-containing proteins Associated With Kleisins (HAWKs), named Ycs4 and Ycg1 in yeast. Ysc4 stably binds to a central region of Brn1 proximal to the neck (HAWK^neck^), whereas Ycg1 binds to a central region proximal to the cap (HAWK^cap^)(Hassler et al., 2018; Uhlmann, 2016; Yatskevich et al., 2019).

Biochemical and structural analyses have exposed a variety of conformational states and DNA interacting modes of SMC complexes (Figure S1B). The two ATPase heads engage with each other upon binding a pair of ATP molecules between them (Lammens et al., 2004). The orientation of the heads in the engaged state spreads apart the coiled-coil arms (Hassler et al., 2019; Vazquez Nunez et al., 2021). Upon ATP hydrolysis, the heads disengage and rotate allowing the coiled-coil SMC arms to align into a rod-shaped structure (Diebold-Durand et al., 2017; Soh et al., 2015). Both in the spreaded and aligned conformations, the SMC arms can bend at an elbow region, allowing the hinge domain to reach the vicinity of the ATPase heads (Burmann et al., 2019; Eeftens et al., 2016; Ryu et al., 2020). The HAWKs subunits are also highly flexible and dynamic within the complex. In yeast condensin, the Ycs4-Brn1 module interacts with the two head domains, both in the apo and engaged states; whereas the Ycg1-Brn1 module is peripheral and more mobile although it can also interact with the other condensin subunits (Hassler et al., 2019; Lee et al., 2020; Lee et al., 2022). Early studies identified the hinge as a DNA binding module, which presents affinity for single-and double-stranded DNA (Griese et al., 2010; Hirano and Hirano, 2006). Another DNA binding module is the kleisin-HAWK^cap^ complex, which secures the DNA with a kleisin belt (Kschonsak et al., 2017; Li et al., 2018). Lastly, SMC complexes form a central DNA clamping module upon ATP binding, in which DNA is hold between the engaged heads and the kleisin-HAWK^neck^ complex (Burmann et al., 2021; Higashi et al., 2020; Lee et al., 2022; Shaltiel et al., 2021; Shi et al., 2020). In addition, DNA can be found topologically or pseudo-topologically entrapped inside the tripartite ring structure (Cuylen et al., 2011; Haering et al., 2008; Ivanov and Nasmyth, 2005; Murayama and Uhlmann, 2014) or in other kleisin-encircled chambers as in the kleisin-HAWK^cap^ complex (Collier et al., 2020; Kschonsak et al., 2017; Shaltiel et al., 2021).

Numerous models are currently postulated for the loop extrusion mechanism of SMC complexes. The ‘‘walking’’ and ‘‘inchworm” models propose that the coiled-coils and ATPase heads function like legs that walk or slide along the DNA (Fudenberg et al., 2016; Nichols and Corces, 2018). The ‘‘pumping” or “segment capture” model speculates that a DNA segment bound at the hinge is pushed towards the head domains via the zipping of the coiled-coils (Diebold-Durand et al., 2017; Marko et al., 2019). The ‘‘scrunching’’ and “swing and clamp” models postulate that motions of SMC arms from extended to bent conformations serve to transfer a DNA segment from the hinge to the ATPase heads (Bauer et al., 2021; Ryu et al., 2020). The “brownian ratchet” model posits that a DNA clamped via head engagement is only allowed to slip unidirectionally, aided by the motion of the SMC arms (Higashi et al., 2021). To date, it is unknown which, if any, of these models is correct. However, a common trait of these proposed mechanisms is their large impact on the topology of the interacting DNA, which is either pushed, pulled or bent. In this respect, earlier *in vitro* studies had revealed that condensin is able to restrain DNA (+) supercoils in an ATP-dependent manner (Kimura and Hirano, 1997, 2000; Kimura et al., 1999; St-Pierre et al., 2009; Takemoto et al., 2006). Since this topological effect required high concentrations and molar ratios of condensin to DNA, its mechanistic significance has not been further investigated. Here, we analysed how condensin alters the topology of the interacting DNA in experimental conditions that sustain DNA loop extrusion (Ganji et al., 2018). Surprisingly, we found that during each round of ATP usage, condensin restrains DNA negative supercoils by constricting a short left-handed loop of DNA, which is not in the extruded loop region. We propose a general mechanistic scheme for how SMC complexes generate DNA translocation steps and extrude DNA loops based on these findings.

## RESULTS

### Catalytic amounts of condensin restrain negative DNA supercoils during ATP usage

The linking number (Lk) of double stranded DNA in a covalently closed domain equals the sum of the DNA twist (Tw or helical winding of the duplex) and the DNA writhe (Wr or non-planar bending of the duplex). Accordingly, ΔLk=ΔTw+ΔWr, meaning that any change of Tw and/or Wr constrained by a DNA binding factor can be revealed by resetting (i.e. relaxing) the Lk of the DNA with a topoisomerase (Figure S2). Following this notion, several studies had shown that, when relaxed DNA plasmids are incubated with condensin and ATP, topoisomerases increase the Lk of the DNA (Kimura and Hirano, 1997, 2000; Kimura et al., 1999; St-Pierre et al., 2009; Takemoto et al., 2006). These observations lead to the conclusion that condensin restrained DNA (+) supercoils. However, constraining of such (+) supercoils (or more precisely, positive ΔLk values) required high concentrations (>50 nM) and molar ratios of condensin to DNA (>1 complex/100 bp). Hence, we asked whether low concentrations and molar ratios of condensin, as those supporting DNA loop extrusion, could also produce measurable ΔTw and ΔWr deformations in the DNA. To this end, we incubated different amounts of the budding yeast condensin (Figure S3) with a relaxed DNA plasmid (4.3 kb) in presence of vaccinia virus topoisomerase I (Topo I). To determine ΔLk changes accurately, we examined the resulting distribution ladders of Lk topoisomers in 1D or 2D agarose gel electrophoreses containing calculated amounts of chloroquine (Figure S4).

We first tested high concentrations and molar ratios of condensin to DNA (Figure 1A and 1B). In the absence of ATP, Topo I did not significantly alter the Lk of the relaxed DNA (R), even when mixed with high concentrations (240 nM) and molar ratios (80:1) of condensin. However, upon addition of ATP, Topo I increased the Lk of the plasmid proportionally to the amount of condensin (Figure 1A and 1C), in agreement with the restraining of (+) supercoils observed in earlier studies (Kimura and Hirano, 1997, 2000; Kimura et al., 1999; St-Pierre et al., 2009; Takemoto et al., 2006). We determined that the ΔLk restrained per condensin was about +0.15 (+6/40), which denoted that each holo-complex might be stabilizing a slight overtwisting (ΔTw≈+0.15) or right-handed bending (ΔWr≈+0.15) of the DNA (Figure S5) (Vologodskii and Cozzarelli, 1994).

**Figure 1.**
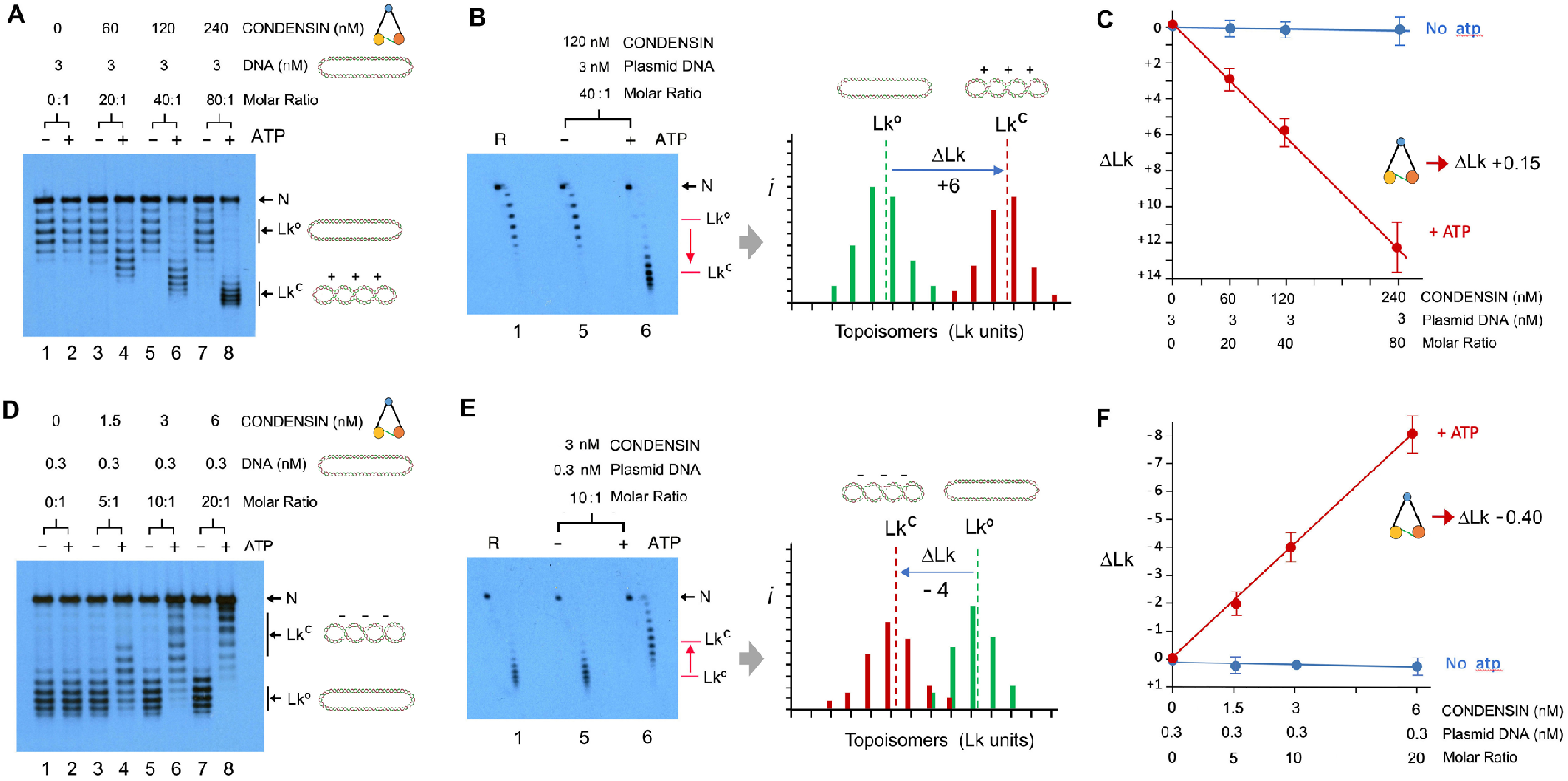
Effect of condensin concentration to restrain DNA supercoils. (A) Relaxed DNA (3 nM), condensin (0, 60, 120, 240 nM) and Topo I (1 unit) were mixed in 25 mM Tris-HCl pH 7.5, 25 mM NaCl and 5 mM MgCl_2_, 1 mM DTT. One half of each mixture was supplemented with ATP (1 mM) and incubations proceeded at 30ºC for 30 min. DNA electrophoresis contained 0.1 µg/ml chloroquine. Reaction settings and electrophoretic analyses are detailed in the methods. In all gels, N denotes nicked circles; Lkº is the input Lk distribution of relaxed DNA; and Lk^C^ the Lk distribution restrained by condensin. (B) 2D-gel of the preceding samples in lane 1 (relaxed DNA, R), and lanes 5 and 6 (condensin 120 nM ± ATP). Electrophoresis contained 0.1 µg/ml and 1 µg/ml chloroquine respectively in the first and second dimension. The histogram shows the relative intensity (*i*) of individual topoisomers of the Lk distributions resolved in lane 1 (green) and lane 6 (red). Lkº and Lk^C^ denote the midpoint of each *Lk* distribution, and ΔLk the difference (Lk units) between them. (C) Plot of positive ΔLk values restrained by indicated concentrations and molar ratios of DNA and condensin with and without ATP (mean ±SD from three independent experiments). (D) Experiment conducted as in A, but reducing the concentration of DNA (0.3 nM) and condensin (0, 1.5, 3, 6 nM). DNA electrophoresis contained 0.4 µg/ml chloroquine. (E) 2D-gel of the preceding sample in lane 1(relaxed DNA, R), and lanes 5 and 6 (condensin 3 nM ± ATP). Electrophoresis contained 0.4 µg/ml and 1 µg/ml chloroquine respectively in the first and second dimension. The histogram shows relative Lk intensities (*i*) of lanes 1 and 6, indicating Lkº, Lk^C^ and ΔLk. (F) Plot of negative ΔLk values restrained by indicated concentrations and molar ratios of DNA and condensin with and without ATP (mean ±SD from three independent experiments).

We next tested reducing the concentration and molar ratios of condensin to DNA over 10-fold, thus mimicking the reaction settings that support DNA loop extrusion (Ganji et al., 2018; Kim et al., 2019). In these conditions, condensin activity did no longer restrain (+) supercoils. Surprisingly, it instead restrained (-) supercoils (Figure 1D and 1E). Namely, Topo I reduced the Lk of the DNA, producing a ΔLk of about -0.4 per condensin complex in an ATP dependent manner (Figure 1F). This topological effect correlated with the amount of condensin and was observable with molar ratios as little as one condensin complex per plasmid (Figure S6). These findings denoted that, upon ATP usage, each complex restrained a significant untwisting (ΔTw≈-0.4) or a left-handed turn (ΔWr≈-0.4) in the DNA (Figure S5). As in the case of DNA loop extrusion, restraining of (-) supercoils was optimal in low or moderate salt buffers (25 to 100 mM NaCl/KCl) containing divalent cations (1 to 5 mM MgCl_2_) (Figure S7). Restraining of (-) supercoils was robust within physiological ranges of pH (7.5) and temperature (15° to 45°C) (Figure S8).

### Negative supercoils remain constrained by condensin after ATP hydrolysis

Condensin restraining of DNA (-) supercoils occurred quickly following ATP addition and soon reached a plateau (<10 min) irrespectively of the initial concentration of ATP (0.1 to 1 mM) (Figure 2A and 2B). Conversely, incubation of condensin and DNA in presence of AMPPNP, a non-hydrolysable ATP analog, did not produce any significant change in the topology of DNA. Yet, preincubation of condensin and DNA with AMPPNP precluded the effect of ATP subsequently added to the reactions (Figure 2C). Therefore, restraining of (-) supercoils required the hydrolysis of bound ATP.

**Figure 2.**
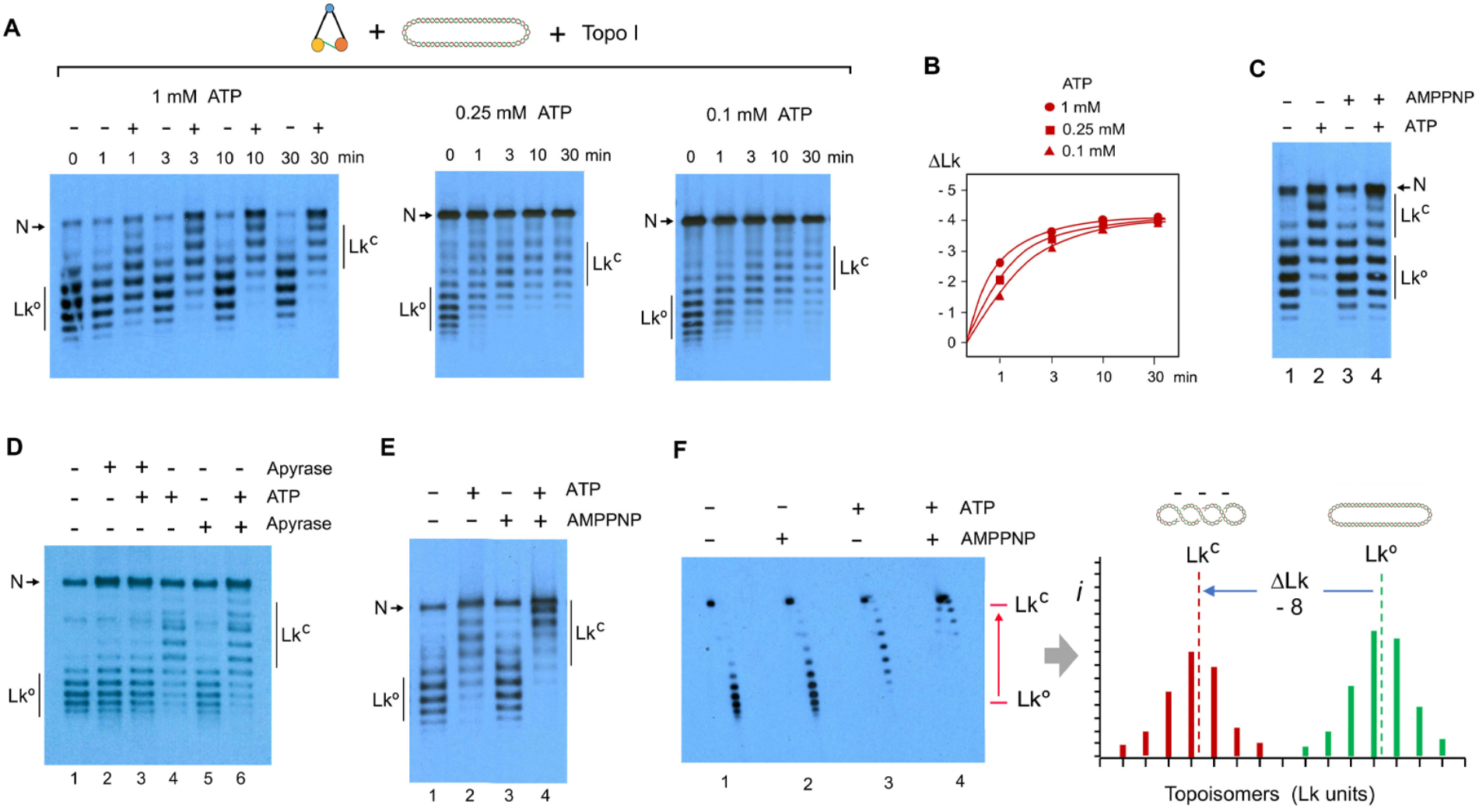
Role of ATP for condensin restraining of negative DNA supercoils. (A) Relaxed DNA (0.3 nM), condensin (3 nM) and Topo I (1 unit) were mixed in 25 mM Tris-HCl pH 7.5, 25 mM NaCl, 5 mM MgCl_2_, 1 mM DTT, and supplemented with different concentrations of ATP (1, 0.25, 0.1 mM). Incubations proceeded at 30ºC for indicated time periods (0 to 30 min). (B) Plot of ΔLk values constrained in the preceding experiment. (C) DNA, condensin and Topo I were mixed as in A. Incubations were at 30ºC without nucleotides for 10 min (lane 1), with ATP 1mM for 10 min (lane 2), AMPPNP 2 mM for 10 min (lane 3), and AMPPNP 2mM for 10 min followed by ATP 1 mM for 10 min (lane 4). (D) DNA, condensin and Topo I were mixed as in A. Incubations were at 30ºC without nucleotides for 10 min (lane 1), with Apyrase for 10 min (lane 2), Apyrase and ATP 1mM for 10 min (lane 3), ATP 1mM for 10 min (lane 4), no nucleotide for 10 min followed by Apyrase for 60 min (lane 5), ATP 1mM for 10 min followed by Apyrase for 60 min (lane 6). (E) DNA, condensin and Topo I were mixed as in A. Incubations were at 30ºC without nucleotides for 20 min (lane 1), with ATP 1mM for 20 min (lane 2), AMPPNP 2 mM for 20 min (lane 3), ATP 1mM for 10 min followed by AMPPNP 2 mM for 10 min (lane 4). (F) 2D-gel of previous samples (lanes 1 to 4) and histogram of Lk intensities (*i*) of lanes 1 and 4, indicating Lkº, Lk^C^ and ΔLk. Electrophoresis in A, C, D, E were as in Figure 1D. Electrophoresis in F was as in Figure 1E.

To test whether restraining of (-) supercoils relied on continuous cycles of ATP usage, we incubated condensin and DNA in the presence of ATP and, afterwards, we added apyrase or alkaline phosphatase to exhaust the ATP. Both ATP hydrolases produced similar results (Figure 2D and S9). As expected, condensin did not restrain (-) supercoils when the ATP hydrolases were added at the beginning of the incubations. However, when the hydrolases were added after 10 min of ATP usage, the (-) supercoils constrained by condensin persisted during extended time periods (60 min). Therefore, continuous use of ATP was not necessary to maintain DNA (-) supercoils restrained.

To further assess that restraining of (-) supercoils did not require continuous cycles of ATP hydrolysis, we incubated condensin and DNA with ATP for 10 min and then we added AMPPNP to quench ATP usage. Surprisingly, such addition of AMPPNP increased by two-fold the amount of (-) supercoils restrained by condensin (Figure 2E). Namely, each condensin complex restrained a ΔLk of about -0.8 (Figure 2F). This observation indicated that, only following the initial cycles of ATP hydrolysis, nucleotide binding produces a conformation that further enhances the restraining of (-) supercoils. Such ΔLk of -0.8 could denote the untwisting of nearly one helical turn of DNA (ΔTw≈ -0.8) or the stabilization of a compact left-handed coil of DNA (ΔWr≈ -0.8) (Figure S5).

### Restraining of supercoils correlates with the stability of condensin-DNA complexes

Since condensin-DNA interactions must be dynamic, we examined the stability of DNA (-) supercoils restrained by condensin. To this end, we tested the interaction of condensin with plasmid DNA in the presence of a molar excess of single- or double-stranded DNA oligonucleotides (ss- or ds-oligos). First, we incubated condensin (3 nM), plasmid (0.3 nM) and Topo I in low salt buffer (25 mM). Afterwards, we added competitor DNAs (100 or 500 nM) and, lastly, ATP. Ds-oligos completely abolished the capacity of condensin to restrain (-) supercoils, whereas ss-oligos produced a lesser effect (Figure 3A). Importantly, oligonucleotides did not affect Topo I activity (Figure S10). Therefore, prior to ATP usage, condensin-DNA interactions must be weak since they were overtaken by competitor DNAs. However, when we incubated condensin, plasmid DNA and Topo I in presence of ATP for 10 min and then added the competitor DNAs, neither ds-nor ss-oligos affected the capacity of condensin to maintain (-) supercoils constrained (Figure 3B). Therefore, condensin-DNA interactions restraining (-) supercoils are either enduring or are reinstated very quickly during each round of ATP usage.

**Figure 3.**
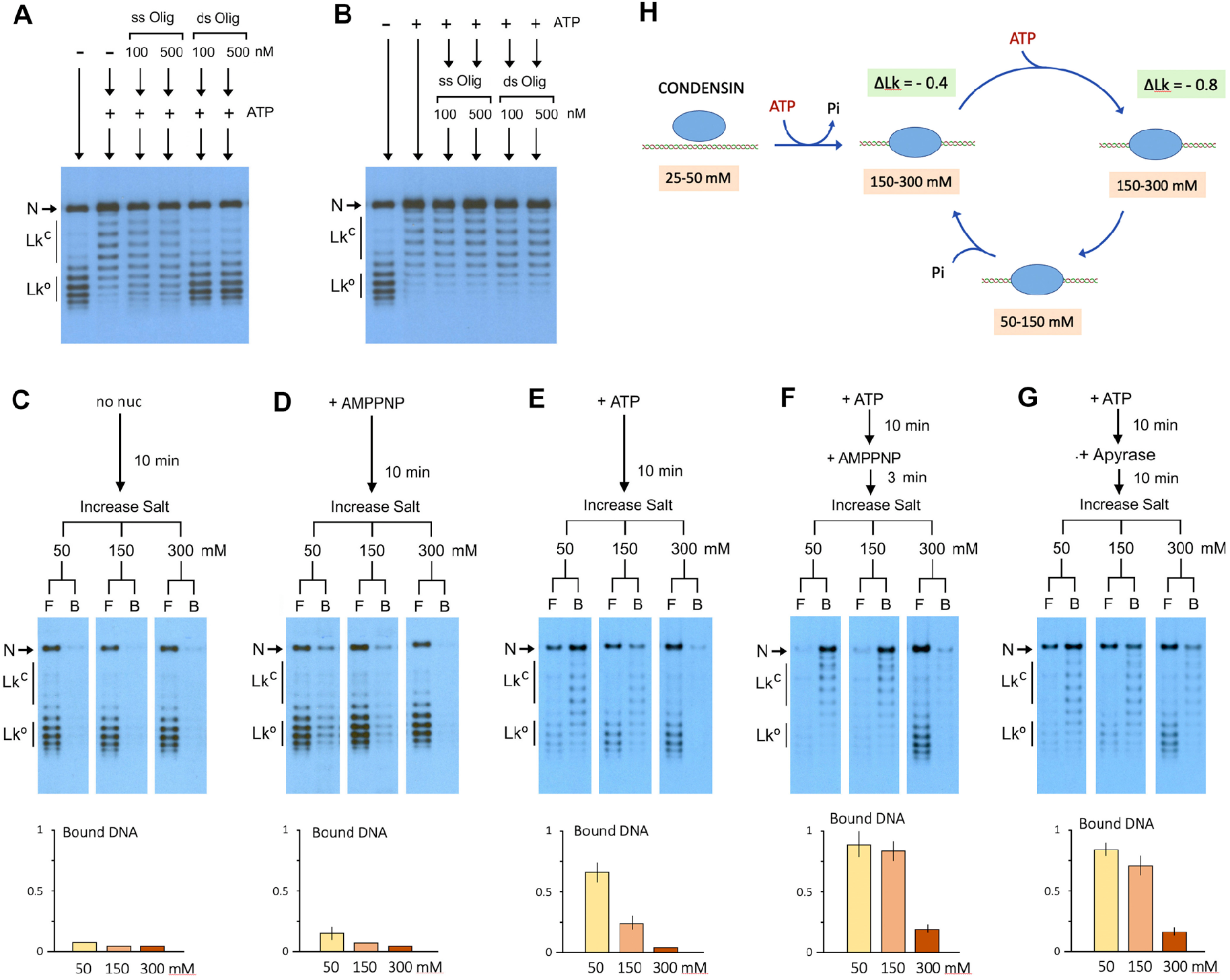
Stability of condensin-DNA complexes during the restraining of DNA supercoils. (A) Relaxed DNA (0.3 nM), condensin (3 nM) and Topo I (1 unit) were incubated in 25 mM Tris-HCl pH 7.5, 25 mM NaCl, 5 mM MgCl_2_, 1 mM DTT. Next, ss-oligos or ds-oligos (100 and 500 nM) were added to the mixtures. Following incubation for 10 min at 30ºC, ATP (1 mM) was added and incubations continued for 10 min. (B) Experiment as in A, but first adding the ATP for 10 min and afterwards the oligonucleotides for 10 min. (C) Relaxed DNA (0.3 nM), condensin (3 nM) and Topo I (1 unit) were mixed in 25 mM Tris-HCl pH 7.5, 25 mM NaCl, 5 mM MgCl_2_, 0.01% Tween-20, 10 mM imidazol. Following incubation at 30ºC for 10 min, reactions were split into thirds to which NaCl concentration was raised to 50, 150 and 300 mM. Salt-resistant condensin-DNA complexes were immobilized to His-Tag magnetic beads and the fractions of free (F) and bound (B) DNA recovered. The bottom plot shows DNA fractions of salt-resistant complexes. (D) As in C, but containing AMPPNP (2 mM). (E) As in C, but containing ATP (1 mM). (F) As in C, but containing ATP (1 mM) for 10 min followed by AMPPNP (2 mM) for 3 min. (G) As in C, but containing ATP (1 mM) for 10 min followed by Apyrase for 10 min. Electrophoresis in A-G were done as in Figure 1D. (H) Condensin-DNA conformation stages inferred from restrained ΔLk values and salt-resistance during ATP usage. See main text for descriptions.

To further assess the stability of condensin-DNA interactions, we next examined the ability of condensin to maintain (-) supercoils restrained under different ionic strength conditions. We incubated condensin, plasmid and Topo I in a low salt buffer (NaCl 25 mM) in the absence or presence of nucleotides. Following the incubations, we increased the salt concentration to 50, 150, 300 mM; and then immobilized condensin to magnetic beads such that we could recover the fractions of free (F) and condensin-bound DNA (B). In the absence of ATP (Figure 3C) or presence of AMPPNP (Figure 3D), nearly all plasmid molecules were free in 50 mM salt. In contrast, when ATP was present in the reactions (Figure 3E), most plasmids were found in the condensin-bound fraction. However, the fraction of bound plasmids was markedly reduced when the ionic strength was raised to 150 mM NaCl. Next, we examined the reactions in the presence of ATP but subsequently quenched by the addition of AMPPNP (Figure 3F). In this case, most plasmids remained bound to condensin when the salt concentration was increased to 150 mM. Only when the salt was raised to 300 mM, the majority of plasmids appeared in the unbound fraction. Lastly, we examined the reactions initiated in presence of ATP but subsequently exhausted by the addition of Apyrase (Figure 3G). Here again, condensin-DNA complexes resisted 150 mM salt; and only after raising the salt to 300 mM, most plasmids were found to dissociate. Importantly, in all cases, the DNA plasmids that remained bound to condensin presented restrained (-) supercoils. Conversely, when DNA plasmids were dissociated from condensin, their (-) supercoils become unconstrained and therefore relaxed by Topo I.

Altogether, the correlation of ΔLk values with ATP usage (Figure 2) and complex stability (Figure 3), indicate that prior to any round of ATP hydrolysis, condensin-DNA interactions are weak and do not alter DNA topology. Following the initial rounds of ATP hydrolysis, the complex gains stability (resisting 50 to 150 mM salt) and constrains ΔLk≈-0.4. In the ensuing rounds of ATP usage, ATP-binding produces a more stable conformation (resisting 150 to 300 mM salt) that restrains ΔLk -0.8. Upon completion of ATP hydrolysis, the complex remains stable (resisting 150 to 300 mM salt) and restrains ΔLk -0.4, implying that condensin-DNA interactions are transiently weakened in the intermediate stages while undergoing ATP hydrolysis (resisting 50 to 150 mM salt) (Figure 3H).

### Condensin-mediated changes of DNA topology do not span outside the condensin-DNA complex

Our experiments were done using Topo I, which is a type-1B topoisomerase that transiently cleaves one strand of duplex DNA and allows free rotation of the other strand in either direction to release (+) or (-) DNA helical tension (Figure 4A) (Champoux, 2001). Accordingly, when condensin restrained (-) supercoils, Topo I relaxed the compensatory (+) supercoiling (helical tension) and so reduced the Lk of DNA. We then expected that any topoisomerase able to relax (+) supercoiling should also reduce the Lk of DNA in these experiments. To this end, we tested topoisomerase II of *s. cerevisiae* (Topo II), which is a type-2A topoisomerase that passes one segment of duplex DNA through a transient double-strand break produced in another segment in an ATP dependent manner (Figure 4B) (Champoux, 2001). The Topo II mechanism relaxes (+) and (-) DNA supercoiling and can also entangle or disentangle DNA molecules.

**Figure 4.**
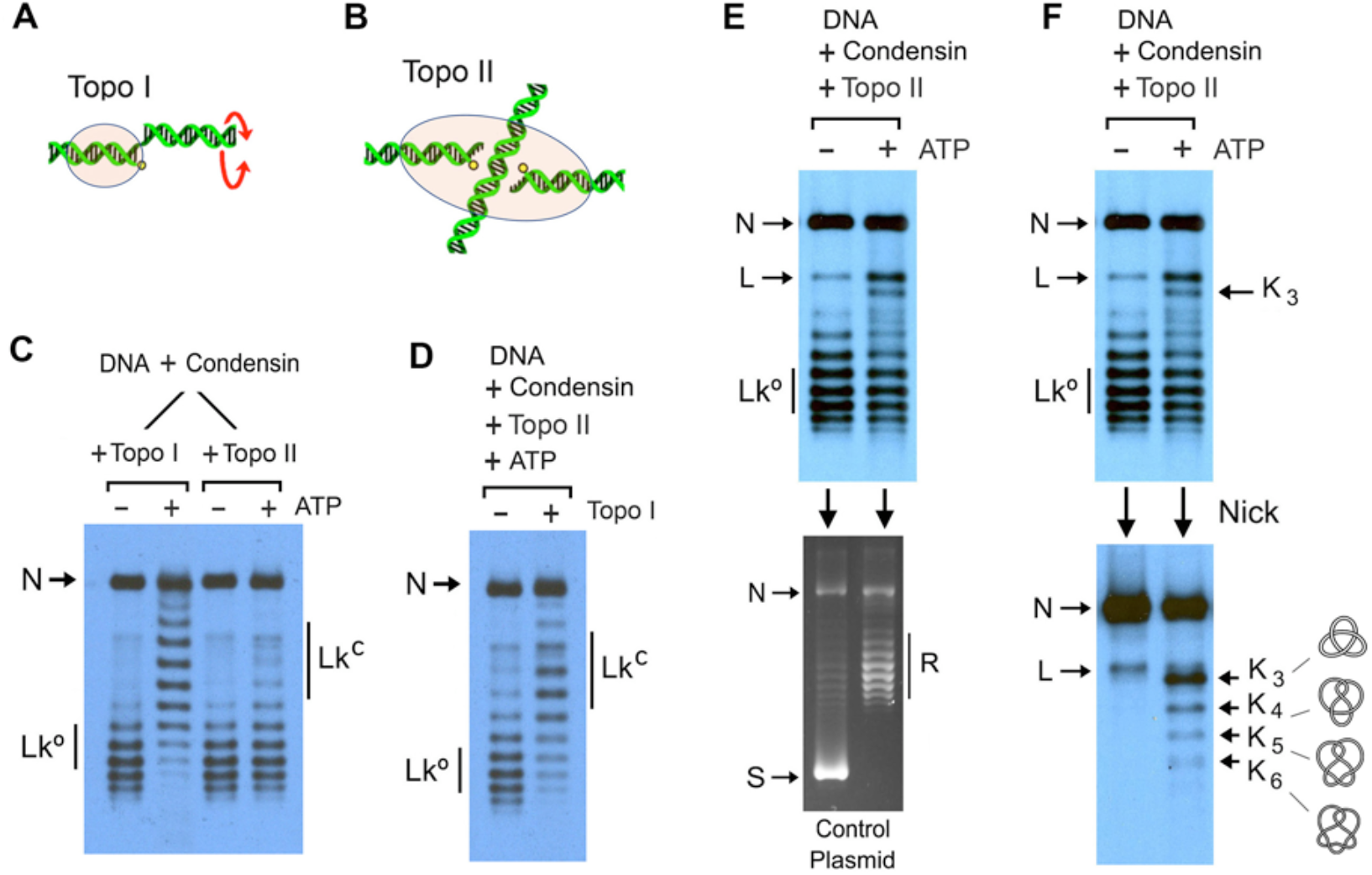
Effect of Topo II activity in the topology of DNA bound to condensin. (A) DNA-strand rotation mechanism of type-1B topoisomerases (Topo I). (B) DNA-cross inversion mechanism of type-2A topoisomerases (Topo II). (C) Relaxed DNA (0.3 nM) and condensin (3 nM) were mixed in 25 mM Tris-HCl pH 7.5, 25 mM NaCl, 5 mM MgCl_2_, 1 mM DTT, either in the presence of Topo I or Topo II. Upon addition of ATP (1 mM), incubations proceeded at 30ºC for 30 min. (D) DNA and condensin, mixed as in C, were supplemented with Topo II and ATP. Following incubation at 30ºC for 10 min, Topo I was added to one half of the mixtures and incubation continued for 10 min. (E) DNA and condensin, mixed as in C, were supplemented with Topo II and with or without ATP. Following incubation at 30ºC for 10 min (top gel), a control (-) supercoiled DNA plasmid (200 ng) was added to one half of the mixtures and incubations continued for 10 min (bottom gel). (F) The other half of the mixtures described in E (same top gel) were nicked with an endonuclease and electrophoresed (bottom gel) to reveal the occurrence of DNA knots. L, linear DNA. K_3_ to K_6_, knots with 3 to 6 irreducible DNA crossings. Electrophoresis in C-F were done as in Figure 1D. The control plasmid (S) relaxed (R) by Topo II in E was stained with Ethidium.

We incubated condensin with relaxed plasmid in the presence of ATP, but included Topo II instead of Topo I in these reactions. Surprisingly, in contrast to Topo I, Topo II did not produce any change in the Lk of the DNA (Figure 4C). We discarded that Topo II could be abrogating the activity of condensin, because the Lk of the plasmid was reduced when Topo I was added to the mixtures already containing Topo II (Figure 4D). We also excluded that condensin could be inhibiting Topo II activity, since Topo II was able to relax supercoiled plasmids subsequently added to the condensin-DNA mixtures (Figure 4E). Moreover, in the mixtures containing Topo II, we observed a burst of DNA knotting in the plasmid (Figure 4F). These knots (K_3_, K_4_, K_5_…) likely resulted from the Topo II-mediated entanglement of intramolecular DNA loops extruded by condensin and, therefore, corroborated that both Topo II and condensin were active in the assay. Therefore, the incapacity of Topo II to reduce the Lk of the DNA indicated that, when condensin restrains (-) DNA supercoils, the compensatory (+) supercoiling is confined within the condensin-DNA complex. Consequently, the restrained (-) supercoils and the compensatory (+) supercoiling must occur within a DNA topological domain bounded by condensin. This topological domain is large enough for the Topo I mechanism to relax the compensatory (+) supercoiling of the DNA (Figure 4A), but not large enough to form and expose a (+) DNA crossover that could be relaxed by Topo II (Figure 4B).

### Condensin does not unwind DNA to restrain negative supercoiling

Condensin could restrain negative ΔLk values either by unwinding the DNA duplex (ΔTw<0) or producing a left-handed turn or loop in the DNA (ΔWr<0) or combining both types of deformations. In either case, we learned that condensin prevents the spreading of compensatory (+) supercoiling to DNA regions outside the complex. With these premises, we envisioned two possible scenarios. One is that condensin uses two DNA binding sites to demarcate a short topological domain of DNA and, afterwards, unwinds or unzips (ΔTw<0) one section of this domain allowing the remaining section to become overtwisted (Figure 5A). The second possibility is that, upon demarcating a short DNA topological domain between two DNA binding sites, these two sites approach each other to bend or loop the DNA in a left-handed manner (ΔWr<0) and, therefore, produce a concomitant overtwisting of the duplex (Figure 5B). Note that, in both scenarios, Topo I (but not Topo II) would be able to relax the overtwisted regions of the DNA domain and so reduce the Lk of the DNA.

**Figure 5.**
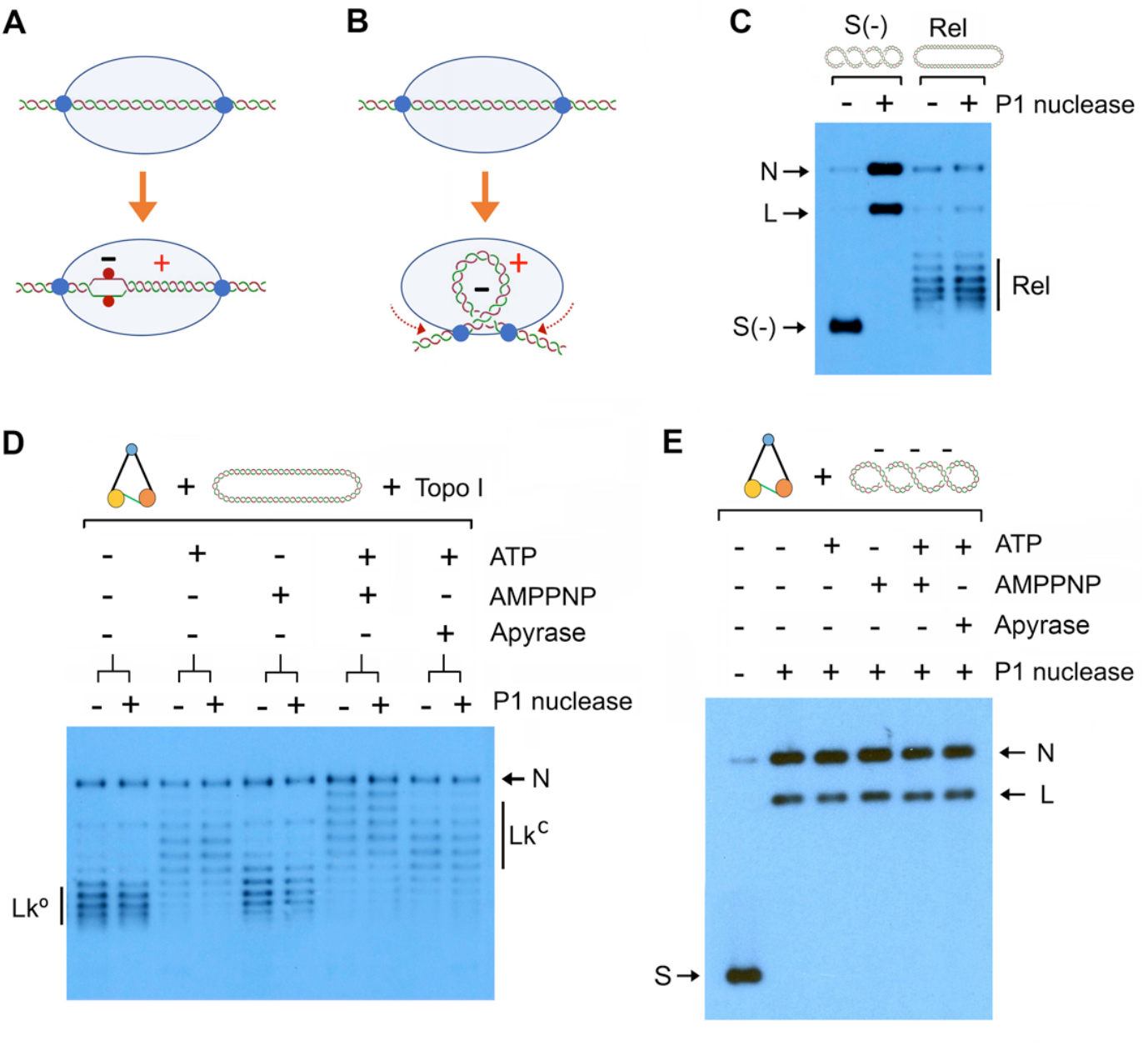
Endonuclease assays to test DNA unwinding by condensin. (A) Two DNA binding sites delimit a short topological domain of DNA and subsequent unwinding (-) of one section is compensated by overtwisting (+) the rest of the domain. (B) Two DNA binding sites delimit a short topological domain of DNA and then approach each other to bend or loop the DNA in a left-handed manner (-), producing concomitant overtwisting of DNA (+). (C) Negatively supercoiled (0.3 nM) or relaxed DNA (0.3 nM) were incubated in the presence or absence of nuclease P1 for 10 min at 30ºC in 25 mM Tris-HCl pH 7.5, 25 mM NaCl, 5 mM MgCl_2_, 1 mM DTT. (D) Relaxed DNA (0.3 nM), condensin (3 nM) and Topo I were mixed in 25 mM Tris-HCl pH 7.5, 25 mM NaCl, 5 mM MgCl_2_, 1 mM DTT. Mixtures were supplemented with/without nuclease P1 and incubated at 30ºC with no nucleotide for 30 min, or ATP 1mM for 30 min, or AMPPNP 2 mM for 30 min, or ATP for 20 min followed by AMPPNP for 10 min, or ATP for 20 min followed by Apyrase for 10 min. (E) Experiment conducted as in D, but using (-) supercoiled DNA and without including Topo I. Following the incubation with nucleotides for 30 min, nuclease P1 was added for 5 min. Electrophoresis in C, D and E were done as in Figure 1D. Supercoiled (S), relaxed (R), nicked (N) and linear DNA signals are indicated.

To test if condensin was unwinding or unzipping the DNA during ATP usage, we employed a single-stranded DNA endonuclease. To this aim, we chose nuclease P1 because this endonuclease does not produce any cleavage (nick) in relaxed DNA but instead is very proficient in nicking negatively supercoiled (i.e., untwisted) DNA (Figure 5C and S11). Thus, we mixed condensin, relaxed DNA plasmid, Topo I and nuclease P1. We then supplemented the mixtures with either ATP, AMPPNP, ATP followed by AMPPNP or ATP followed by Apyrase. Following 30 min incubations, we found that nuclease P1 did not nick at all the DNA plasmids in which condensin was restraining (-) supercoils (Figure 5D). We discarded that condensin could be inhibiting the nuclease by conducting a similar experiment, in which input DNA was negatively supercoiled and Topo I was not included in the reactions. In this experiment, the nuclease was added at the end of the incubations and rapidly nicked all the plasmids (Figure 5E). Therefore, we concluded that condensin restrains (-) supercoils by stabilizing a left-handed DNA turn or loop (Figure S5), rather than by unwinding DNA.

### DNA supercoiling energy alone cannot drive condensin to restrain supercoils

In our experiments, condensin restrained (-) supercoils in relaxed DNA plasmids while Topo I relaxed the compensatory (+) supercoiling thus reducing the Lk of the DNA. However, we also investigated the capacity of condensin to restrain (-) supercoils in a different experimental setting (Figure 6A). Namely, we incubated condensin with a plasmid that was already (-) supercoiled and added Topo I to relax the DNA only at the end of the incubation. In the absence of nucleotides or presence of AMPPNP, Topo I relaxed all the (-) supercoils of the input plasmid (Figure 6B). However, in the mixtures incubated with ATP, ATP followed by AMPPNP or ATP followed by Apyrase, Topo I did not relax all the (-) supercoils, indicating that some had been constrained by condensin (Figure 6B). Importantly, the ΔLk produced by relaxing free (-) supercoils was the same as produced by relaxing the compensatory (+) supercoils when input DNA was already relaxed (Figure 6C).

**Figure 6.**
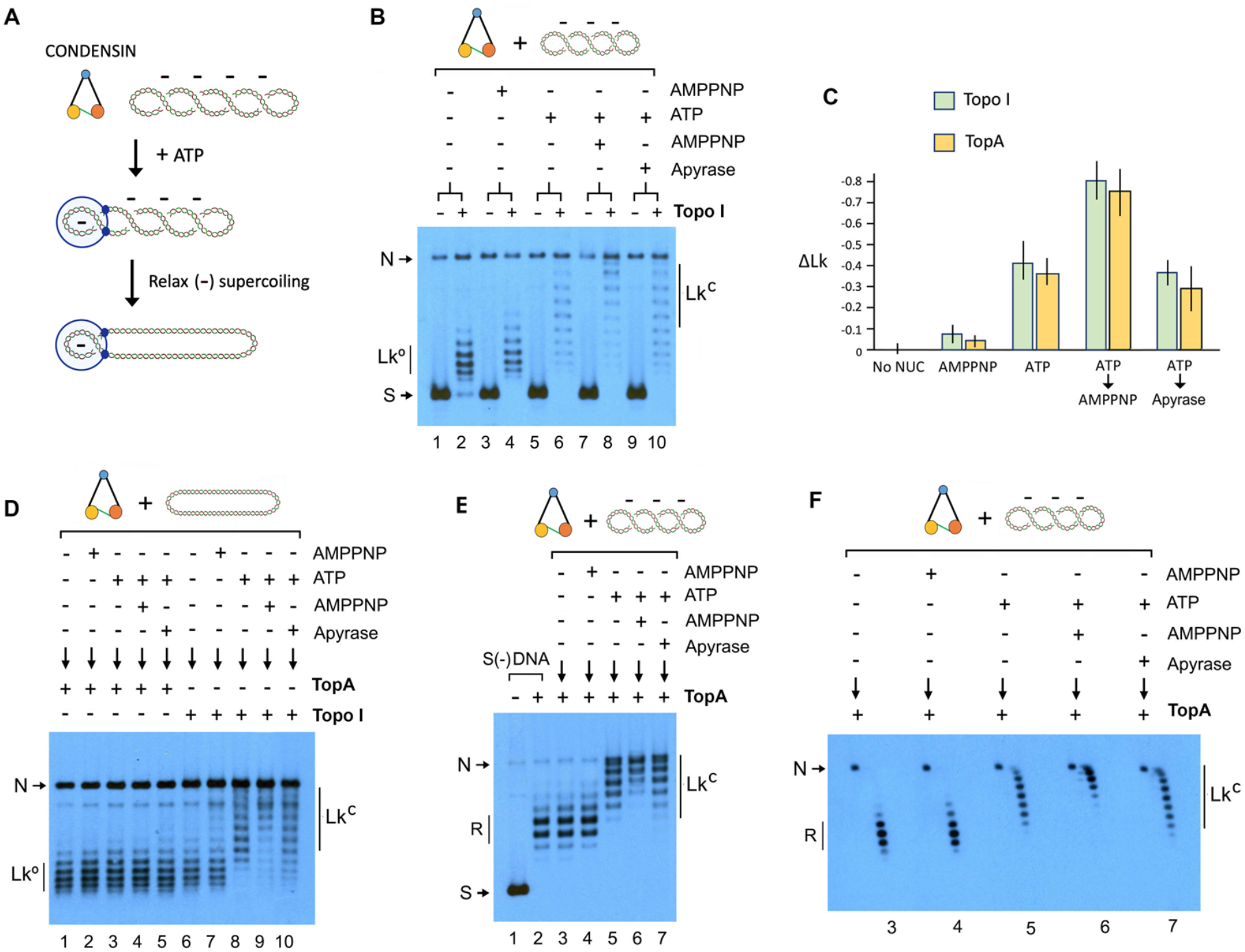
DNA topology restrained by condensin in negatively supercoiled DNA. (A) Condensin ATP-dependent stabilization of a pre-existing (-) supercoil and subsequent relaxation of unconstrained (-) supercoils. (B) (-) supercoiled DNA (0.3 nM) was mixed with condensin (3 nM) in 25 mM Tris-HCl pH 7.5, 25 mM NaCl, 5 mM MgCl_2_, 1 mM DTT. Mixtures were incubated at 30ºC with no nucleotide for 10 min (lanes 1, 2), AMPPNP 2 mM for 10 min (lanes 3, 4), ATP 1mM for 10 min (lanes 5, 6), ATP for 10 min followed by AMPPNP for 10 min (lanes 7, 8), ATP for 10 min followed by Apyrase for 10 min (lanes 9, 10). Topo I was later added to one half of each reaction (even lanes), and incubations continued for 10 min. (C) ΔLk values (mean ±SD from three independent experiments) restrained by condensin when unconstrained (-) supercoils were relaxed by Topo I or TopA. (D) Experiment conducted as in B but using relaxed DNA (0.3 nM). Following incubation in absence or presence of indicated nucleotides, TopA was added to one half of each reaction (lanes 1-5) and Topo I to the other half (lanes 6-10). Incubations continued for additional 10 min. (E) Experiment conducted as in B but, following the incubations with condensin, TopA was added (instead of Topo I) to relax unconstrained (-) supercoils. Lanes 1 and 2 show the (-) supercoiled DNA before and after relaxation with TopA (no condensin added). (F) 2D-gel of samples shown in E (lanes 3 to 7). Electrophoresis in B, D, E were done as in Figure 1D. Electrophoresis in F was done as in Figure 1E.

Since in the above experiment Topo I relaxed (-) supercoils instead of (+) ones, we anticipated that E.coli topoisomerase I (TopA), a type-1A topoisomerase that relaxes (-) but not (+) DNA supercoiling (Champoux, 2001), could also reveal the (-) supercoils restrained by condensin. To test this notion, we first incubated condensin with the relaxed plasmid in the absence or presence of nucleotides and included TopA instead of Topo I. As expected, since TopA cannot relax the compensatory (+) supercoiling, the Lk of the DNA did not change in these reactions (Figure 6D). Next, we incubated condensin with the (-) supercoiled plasmid and added TopA at the end of the incubations (Figure 6E and 6F). In the absence of nucleotides or just the presence of AMPPNP, TopA relaxed the (-) supercoiled plasmid to the same extent observed in relaxation reactions lacking condensin (Figure 6E). However, when we used ATP, ATP followed by AMPPNP or ATP followed by apyrase, TopA relaxed a lesser number of (-) supercoils since part of them had been constrained by condensin (Figure 6E and 6F). Critically, the deficit of (-) supercoils not relaxed by TopA was similar to that produced by Topo I (Figure 6C). These results corroborated that condensin requires ATP to restrain (-) supercoils, irrespective of the input DNA being relaxed or already negatively supercoiled, and that such restrain is maximal during ATP binding.

## DISCUSION

Before the DNA loop extrusion activity of SMC complexes came to light, the capacity of condensin to restrain DNA (+) supercoils was postulated as a mechanism that compacted mitotic chromosomes (Hirano, 2014). However, here we showed that such restraint of (+) supercoils only occurs by using high concentrations or molar ratios of condensin to DNA (i.e. >40 condensins per 4.3 kb plasmid or >1 condensin/100 bp) (Figure 1). When DNA is mixed with such an excess of condensin, far from physiological ratios of about one condensin per 10 Kb (Wang et al., 2005), extrusion of DNA loops is impracticable (Kim et al., 2020). Therefore, in these experimental conditions, the small degree of (+) supercoiling constrained per condensin complex (ΔLk=+0.15) could be the result of unproductive ATP-driven motions of stacked condensins.

By reducing the concentration and molar ratios of condensin to one or few complexes per plasmid DNA, as in current DNA loop extrusion assays, we uncovered that condensin activity restrains (-) supercoils or, more precisely, it restrains negative ΔLk values from -0.4 up to -0.8 per complex (Figures 1 and 2). These changes of DNA topology are coupled to ATP usage in two ways. Following initial cycles of ATP hydrolysis, condensin restrains a ΔLk of -0.4 and this capacity persists irrespective of further ATP consumption. However, in the ensuing cycles of ATP usage, the restrained ΔLk increases to -0.8 upon ATP binding and resets to -0.4 upon completion of ATP hydrolysis (Figure 2 and 3). These findings are consistent with the dual role of the ATPase activity of SMC complexes. Namely, ATP hydrolysis is initially necessary to load the complex onto DNA (Arumugam et al., 2003; Murayama and Uhlmann, 2014; Wilhelm et al., 2015) and the ensuing cycles of ATP consumption allow its translocation along the DNA (Davidson et al., 2019; Ganji et al., 2018; Terakawa et al., 2017). Accordingly, the loaded complex constrains ΔLk of -0.4, but this value raises to -0.8 every time the ATPase heads are engaged via nucleotide binding. These digits imply that the ΔLk of -0.4 restrained during ATP usage is an average value of distinct conformations of the active complex, which include the ATP bound conformation that restrains ΔLk of -0.8, the intermediate stages ongoing ATP hydrolysis, and the post-hydrolysis conformation that restrains ΔLk of -0.4.

Previous studies had denoted the capacity of SMC complexes to interact with ssDNA (Hirano and Hirano, 1998; Sakai et al., 2003). In particular, the hinge domain has more affinity for ssDNA than dsDNA (Griese et al., 2010; Uchiyama et al., 2015); and cohesin is able to interact simultaneously with ds- and ss-DNA molecules (Murayama et al., 2018). These observations pointed to the possibility that condensin restrained negative ΔLk values by unwinding the DNA (ΔTw<0). However, our results indicate that condensin does not produce an unwound region of DNA during its ATP cycle (Figure 5). It seems then unlikely that condensin activity is untwisting nearly a half helical turn (ΔLk -0.4) or near a full helical turn (ΔLk -0.8) of DNA, unless such unwound region of DNA is somehow protected from the nuclease attack. The alternative way for condensin to constrain negative ΔLk values is by stabilizing a left-handed bend or loop of DNA (ΔWr<0). This prospect is more likely than DNA untwisting because SMC complexes necessarily bend the DNA template, in order to start the extrusion process. Moreover, reiterated cycles of DNA bending might occur during loop extrusion. In this respect, experiments using single-molecule magnetic tweezers have shown that condensin (Eeftens et al., 2017) and *Smc5/6* (Gutierrez-Escribano et al., 2020) are able to “lock” DNA plectonemes by embracing their loop stem in an ATP-dependent manner. A cryo EM structure of the bacterial SMC complex (MukBEF) has revealed interactions with two DNA segments with a crossing angle as that produced by a left-handed coil (Burmann et al., 2021). In line with these observations, our results allow to conclude that the negative ΔLk values constrained by condensin are consequent to left-handed bending or looping of DNA.

Another insight of our study is that condensin not only restrains DNA (-) supercoiling, but also impedes the compensatory (+) supercoiling to escape outside the condensin-DNA complex (Figure 4). This observation implies that all these DNA deformations occur within a DNA domain that is enclosed by condensin-DNA interactions. This scenario is easily envisaged if two separated DNA binding sites delimit a DNA topological domain and, afterwards, these sites move towards each other in a way that deforms the enclosed DNA into a left-handed loop. As long as these two DNA binding sites preclude axial rotation of the duplex, the compensatory (+) supercoiling would remain within the formed loop (Figure 5B). Our results also indicate that the compensatory (+) supercoiling occurs in a large enough domain to be relaxed by Topo I, but not large enough to expose a (+) DNA crossover that could be relaxed by Topo II. Therefore, such domain cannot be the extruded loop, which reaches thousands of bp in length. Moreover, DNA loops extruded by SMC complexes are not supercoiled (Davidson et al., 2019; Ganji et al., 2018; Golfier et al., 2020; Kim et al., 2019), which further denotes that Topo I is not relaxing the extruding loop. Accordingly, this short domain relaxable by Topo I must be the segment of DNA that condensin has to restrain during each round of ATP usage to translocate along DNA. In this respect, former analyses of DNA extrusion step sizes of yeast condensin showed values of around 200 nm (Eeftens et al., 2017), but more recent measurements revealed step sizes between 17 and 40 nm (Ryu et al., 2022). These step sizes translate into domains of about 100±50 bp of naked DNA, which could certainly be relaxed by the strand-rotation mechanism of Topo I, but not by the DNA-cross inversion mechanism of Topo II (Figure 4).

The recurrent formation of a short left-handed loop of DNA, topologically bounded by condensin during each round of ATP usage, hint at a general mechanistic scheme of how DNA translocation steps might occur during loop extrusion. This scheme involves three DNA binding modules (the “anchor”, the “mid” and the “grabber”), two DNA loops (the “feeding” loop and the “extruding” loop) and five steps (“load”, “grab”, “pinch”, “merge” and “reset”) (Figure 7A). To start, initial cycles of ATP hydrolysis load the condensin onto DNA occupying the anchor and the mid sites, such that a first domain of DNA is delimited. The loaded complex enables the grabber to seize an additional DNA site and thus enclose a second DNA domain. At this stage and prior ATP binding, the DNA can be deformed by fluctuating movements of the DNA-binding modules. In particular, the grabber and the mid frequently move towards each other producing a recurrent bending of the enclosed DNA. This oscillation is ended upon ATP binding, which fixes the grabber close to the mid and, therefore, pinches the enclosed DNA into a tight loop, which is the feeding loop. Sequential hydrolysis of ATP drives the merging process, which involves two operations. First, the mid site releases its DNA. As a result, the feeding loop is no longer constrained and merges with the DNA domain formerly bounded by the anchor site. Second, the grabber transfers its DNA to the mid site. As a result, the anchor and the mid site enclose an enlarged DNA domain, which is the extruded loop. The reset step enables the grabber to seize a new segment of DNA and repeat the cycle.

**Figure 7.**
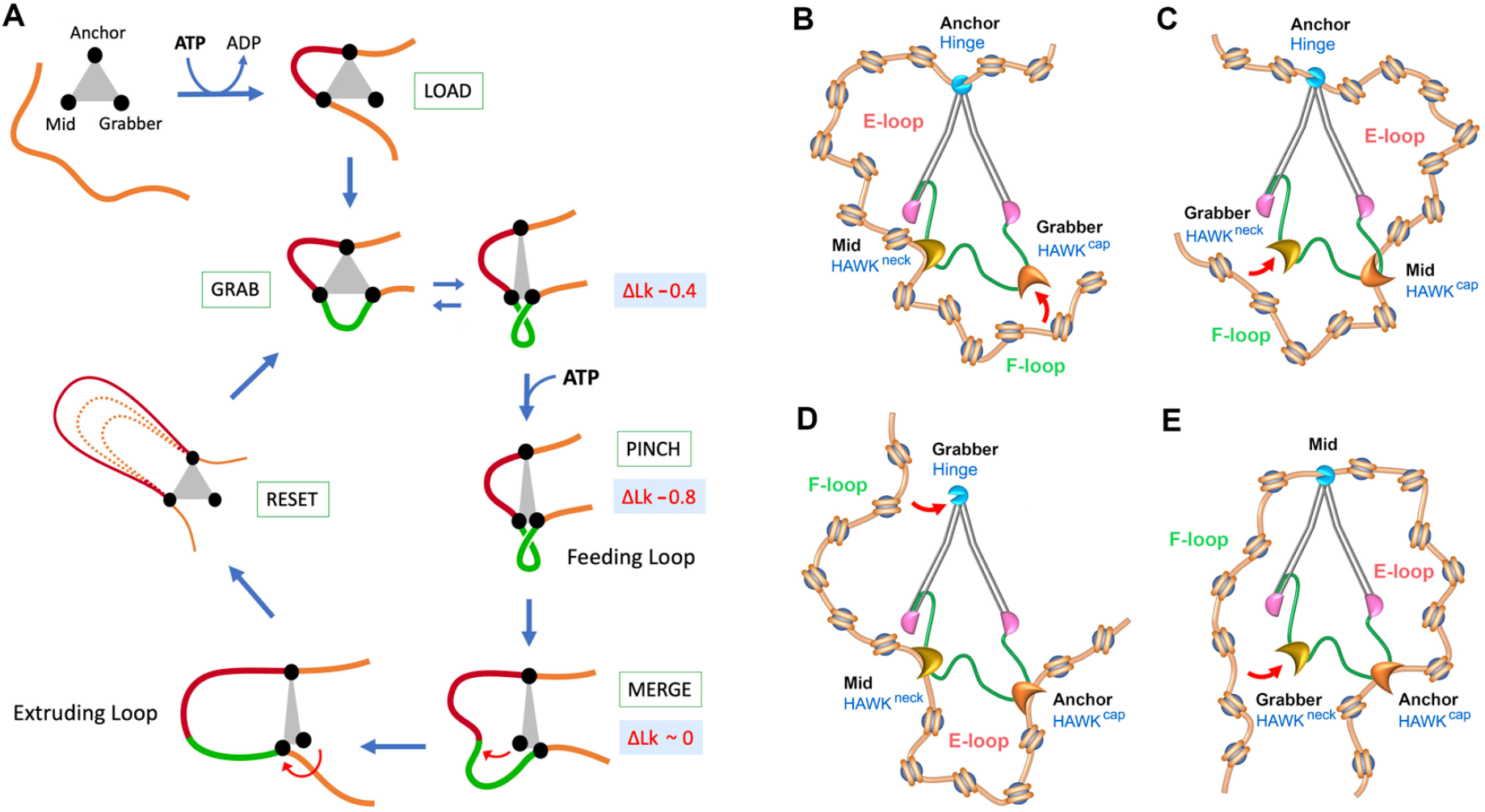
Integration of DNA topology into a mechanistic model of loop extrusion. (A) Mechanistic scheme of DNA loop extrusion by an SMC complex (grey) with three DNA binding modules (Anchor, Mid, Grabber). Following the loading process, each round of ATP usage includes four steps (Grab, Pinch, Merge, Reset), which produce a feeding loop (green) and the subsequent enlargement of an extruded loop (red). The ΔLk values restrained during the grabbing, pinching and merging steps are indicated. See main text for details. (B-E) Plausible roles of three DNA binding modules of condensin (Hinge, HAWK^cap^, HAWK^neck^) to perform the pinch and merge mechanism illustrated in (A). The anchor function is assigned to the hinge in A and B; and to the Kleisin-HAWK^cap^ module in C and D. In each model, a nucleosomal fiber interacts with the anchor and the mid sites, which delimit an incipient extruding loop (E-loop). A third interaction with the grabber site (red arrow) will determine the step size by enclosing a potential feeding loop (F-loop). Conformation models of the ensuing pinching, merging and reset steps are illustrated in Figure S12.

The above mechanistic scheme (Figure 7A), for short named as “pinch and merge”, explains the ΔLk changes observed in our study on the basis that the feeding loop produced by the pinching step is a left-handed loop that restrains ΔLk -0.8 (Figure S5). Accordingly, the restrained ΔLk of -0.4 observed during regular cycles of ATP usage is the mean value produced by the pinching step and the other steps. Note that, during the merging process, the topological constrains of the feeding loop are gone allowing the extruded loop to be virtually relaxed. Also note that the ΔLk of -0.4 observed upon completion of ATP hydrolysis would reflect the recurrent bending of DNA produced by the motions of the grabber and mid sites. The pinch and merge scheme also accounts for the changes in the stability of the condensin-DNA complex observed in our study. During the pinching step (ATP-bound conformation) and after the completion of ATP hydrolysis (ATP exhaustion), the complex is stable because the DNA is held by the three DNA binding modules. However, during the merging process (during ATP-hydrolisis), the complex is less stable since the grabber is transfering its DNA to the mid site.

The pinch and merge mechanism also accounts for recent observations that ATP binding triggers the stepping power in cohesin (Bauer et al., 2021) and condensin (Ryu et al., 2022; Shaltiel et al., 2021). Note that the pinching of the feeding loop produces the main pulling stroke on the DNA. Another quality of the pinch and merge mechanism is that it does not involve sliding movements to tranfer the DNA from one binding site to another. This feature provides the crucial capacity of condensin to bypass nucleosomes, other condensin complexes (Kim et al., 2020) and obstacles much larger than the SMC’s dimensions (Pradhan et al., 2021). Related to this, SMC complexes can extrude DNA loops without topological entrapment of DNA inside the tripartite ring (Davidson et al., 2019; Pradhan et al., 2021; Shaltiel et al., 2021). The pinch and merge mechanism does not require such topological entrappment, not even a pseudo-topological embracement of the feeding loop or the extruding loop in the lumen of the tripartite ring. However, it is likely that condensin might use permanent (anchor) or transient (mid and grabber) mechanical bonds to secure DNA loop boundaries or facilitate their transfer from one module to another.

Since condensin has three main DNA binding modules, the “pinch and merge” scheme can be mechanistically modelled into six posible combinations depending on the role assigned to each module (Figure S12). A priori, since the HAWK^cap^ module is peripheral and highly mobile, it could suit the role of the grabber or the mid site. Likewise, the transient clamping of the DNA between the engaged heads and the HAWK^neck^ module does not fit the role of an stable anchor. These premises would suggest the hinge region as the anchor site (Figures 7A and 7B). However, recent experimental evidence indicated that the HAWK^cap^ module functions as the anchor site during loop extrusion (Shaltiel et al., 2021). This finding leaves the hinge acting as the grabber and the HAWK^neck^ module as the mid site, or vice-versa (Figures 7C and 7D). Future research on the effect of individual DNA binding modules in the restraining of DNA supercoils could markedly narrow these models towards the definitive SMCs mechanism of loop extrusion. Thus far, our study disclosed that, in addition to the anchor site, two additional sites tightly constrain and deform a short domain of DNA into a negatively supercoiled loop when SMC heads are engaged via ATP binding.

## METHODS

### Enzymes and DNA

Yeast condensin was expressed and purified as reported previously (Lee et al., 2020). Briefly, S. cerevisiae cells were transformed with a pair of 2μ-based high copy plasmids containing pGAL10-YCS4 pGAL1-YCG1 TRP1 and pGAL7-SMC4-StrepII3 pGAL10-SMC2 pGAL1-BRN1-His12-HA3 URA3. Overexpression was induced by addition of galactose to 2%. Cell lysates were cleared by centrifugation, loaded onto a 5-mL HisTrap column (GE Healthcare) and eluted with imidazole. Eluate fractions were incubated with Strep-Tactin Superflow high capacity resin and eluted with desthiobiotin. Eluates were concentrated by ultrafiltration and final purification proceeded by size-exclusion chromatography with a Superose 6 column. Purified condensin, recovered at about 3 µM concentration in 50 mM Tris-HCl pH 7.5, 200 mM NaCl, 1 mM MgCl_2_, 1 mM DTT, 5% glycerol, was snap-frozen and stored at –80°C. Before each experiment, condensin was diluted to 300 nM concentration in 50 mM Tris-HCl pH 7.5, 1 mM EDTA, 200 mM NaCl, 1 mM DTT, 500 µg/mL BSA, 50% Glycerol and kept at -20°C until mixing with DNA.

Topoisomerase I of vaccinia virus (Topo I) was expressed and purified from *E. coli* cells harboring the expression clone pET11vvtop1 (Shuman et al., 1988). We defined 1 unit of Topo I as the amount of enzyme that catalysed the relaxation of 100 ng of negatively supercoiled pBR322 DNA in 5 minutes at 30°C in a reaction volume of 20 μl. Topoisomerase II of *S. cerevisiae* (Topo II) was expressed and purified from yeast cells carrying the expression clone YEpTOP2GAL1 (Worland and Wang, 1989). We defined 1 unit of Topo II as the amount of enzyme that catalysed the relaxation of 100 ng of negatively supercoiled pBR322 DNA in 5 minutes at 30°C in a reaction volume of 20 μl. Additional enzymes were from commercial sources: E.coli Topoisomerase I (TopA) (NEB #M0301S); Nuclease P1 (NEB #M0660S); Alkaline phosphatase (NEB #M0290); Apyrase (NEB #M0398S); Endonuclease BstNB1(NEB #R0607).

To produce a stock of relaxed DNA, 10 µg of negatively supercoiled pBR322 (4,3 Kbp) were pre-incubated at 30°C for 5 min in 100 μl of 25 mM Tris-HCl pH 7.5, 25 mM NaCl, 5 mM MgCl2, 1 mM DTT. 10 units of Topo I were added and the incubations proceeded for 30 min. Reactions were terminated with one volume of 20 mM EDTA and 1% SDS, and extracted twice by phenol-chloroform. DNA was recovered by EtOH precipitation and resuspended in 10 mM Tris-HCl pH 7.5, 1 mM EDTA.

### Reactions of DNA with condensin

Incubations of DNA with condensin were typically done in 20 µL of reaction buffer containing 25 mM Tris-HCl pH 7.5, 1 mM DTT, 25 mM NaCl and 5 mM MgCl_2_, unless some components were modified as indicated in specific experiments. The DNA (relaxed or negatively supercoiled pBR322) was first added at the specified final concentrations (0.3 or 3 nM) followed by condensin at the specified final concentrations (0.3 to 240 nM). Upon preincubation at 30ºC for 5 min, the DNA-condensin mixtures were supplemented with either 1 mM ATP, 2 mM AMPPNP, 1 mM ATP subsequently quenched by 2 mM AMPPNP, or 1 mM ATP subsequently exhausted by 1 unit of Apyrase. Reaction mixtures were also supplemented with either 1 unit of Topo I, 1 unit of Topo II, or 1 unit of TopA when indicated. Reactions proceeded at 30ºC for specified time periods until were terminated by adding 10 µl of 20 mM EDTA, 1% SDS, 30% Glycerol, 0.3 µL of proteinase-K (10 mg/mL) and incubated for 30 min at 50°C. Resulting 30 µL volumes were cooled at room temperature and 15 µL loaded in agarose gels for electrophoresis.

### DNA competition assays

DNA competition assays were done in a 20 µL of reaction buffer by first mixing relaxed pBR322 (0.3 nM), condensin (3 nM) and Topo I. Following 5 min incubation at 30ºC, ss- and ds-oligonucleotides (60 base-pairs) were added at high concentration (100 and 500 nM) and incubations continued for 10 min at 30ºC. Reactions were then supplemented with ATP (1 mM) and further incubated for 10 min. Parallel experiments were conducted by first incubating the condensin-DNA mixtures with ATP (1 mM) for 10 min, and afterwards adding the oligonucleotides and continuing the incubation for 10 min. Reactions were terminated and processed as described above.

### Immobilization of condensin-DNA complexes

Relaxed pBR322 (0.3 nM) condensin (3 nM) and Topo I were mixed in a 60 µL volume of buffer containing 25 mM Tris-HCl pH 7.5, 25 mM NaCl and 5 mM MgCl_2_, 0.01% Tween-20, 10 mM Imidazol. Following a preincubation for 5 min, the mixtures were supplemented with nucleotides and incubations proceeded at 30°C for 10 min. Reaction volumes were then divided into thirds of 20µL, to which NaCl was added to reach concentrations of 50, 150, or 300 mM. Following 5 min incubation, 1 µL of His-Tag magnetic beads (Dynabeads™ Invitrogen #10103D) was added to each tube. After 5 min incubation, reaction tubes were placed on the magnet for 2 min and the supernatant containing free DNA was recovered. To release the DNA immobilized by the his-tagged condensin, the magnetic beads were resuspended in 20 µL of 10 mM Tris-HCl pH 7.5, 1 mM EDTA, 1% SDS, 0.3 µL of proteinase-K (10 mg/mL) and incubated for 10 min at 50°C. The beads were centrifuged and the supernatant recovered. 10 µL of 20 mM EDTA and 30% Glycerol were added to the supernatants. 15 µL of the final volumes were loaded in agarose gels.

### Nuclease P1 digestions

Relaxed pBR322 (0.3 nM), condensin (3 nM), Topo I (1 unit) and Nuclease P1 (10 units) were mixed in a 20 µL volume of reaction buffer. In the reactions that started with negatively supercoiled pBR322, Topo I was omitted. Following a preincubation for 5 min, the reaction mixtures were supplemented with nucleotides. Incubations proceeded at 30°C for indicated time periods until were terminated and processed as described above.

### Electrophoresis of DNA topoisomers and calculation of ΔLk

Topoisomers of pBR322 were electrophoresed in 0.7% (w/v) agarose gels. One-dimensional electrophoreses were carried out at 2.5 V/cm for about 20 h in TBE buffer (89 mM Tris-borate, 2m M EDTA) containing 0.4 µg/ml chloroquine (or as specified). In these conditions, topoisomers around Lkº move ahead of the nicked DNA circles, and topoisomers with Lk values higher than Lkº move faster than Lkº. Two-dimensional electrophoreses were in TBE containing 0.2 µg/ml chloroquine in the first dimension (2.5 V/cm for 18 h, gel top to bottom) and in TBE containing 1 µg/ml chloroquine in the second dimension (5 V/cm for 4 h, gel left to right). In these conditions, topoisomers distribute in an arch, in which Lk values increase clockwise and decrease anti-clockwise. Gels were blot-transferred to a nylon membrane (Amersham Hybond-N+ #RPN203B) and probed with pBR322 DNA sequences labelled with AlkPhos Direct (GE Healthcare® #GERPN3680). Chemiluminescent signals of increasing exposition periods were recorded with a cooled CCD camera (KODAK Gel Logic 1500 Imaging System) or on X-ray films. Lk changes were analysed as described (Segura et al., 2018). Briefly, the midpoint of each Lk distribution, which does not necessarily coincide with the position of one DNA topoisomer, was determined by quantifying with the ImageJ software the relative intensity of non-saturated signals of the individual Lk topoisomers. *ΔLk* was calculated as the distance (Lk units) between the midpoints of the input relaxed DNA distribution (Lkº) and of the Lk distribution restrained by condensin activity (Lk^C^). To observe the knot species produced by Topo II, the reacted DNA samples were nicked with endonuclease BstNB1 and examined in one-dimensional electrophoreses as described for Lk topoisomers.

## ACKNOWLEDGMENTS

Work in the Roca laboratory is supported by the Plan Estatal de Investigación Científica y Técnica of Spain, with grant PID2019-109482GB-I00 to J.R; and research fellowships BES-2016-077806 to S.D. and BES-2012-061167 to J.S. Work in the Aragon laboratory is supported by the Medical Research Council (UKRI MC-A652-5PY00).

## AUTHOR CONTRIBUTIONS

Conceptualization, J.R. and L.A.; Methodology, J.R.; Investigation, J.S., S.D, and B.M-G; Resources, P.G-E. and L.A.; Writing, Visualization, and Supervision, J.R.

## DECLARATION OF INTERESTS

The authors declare no competing interests.

## Supplemental Information

**Figure S1.**
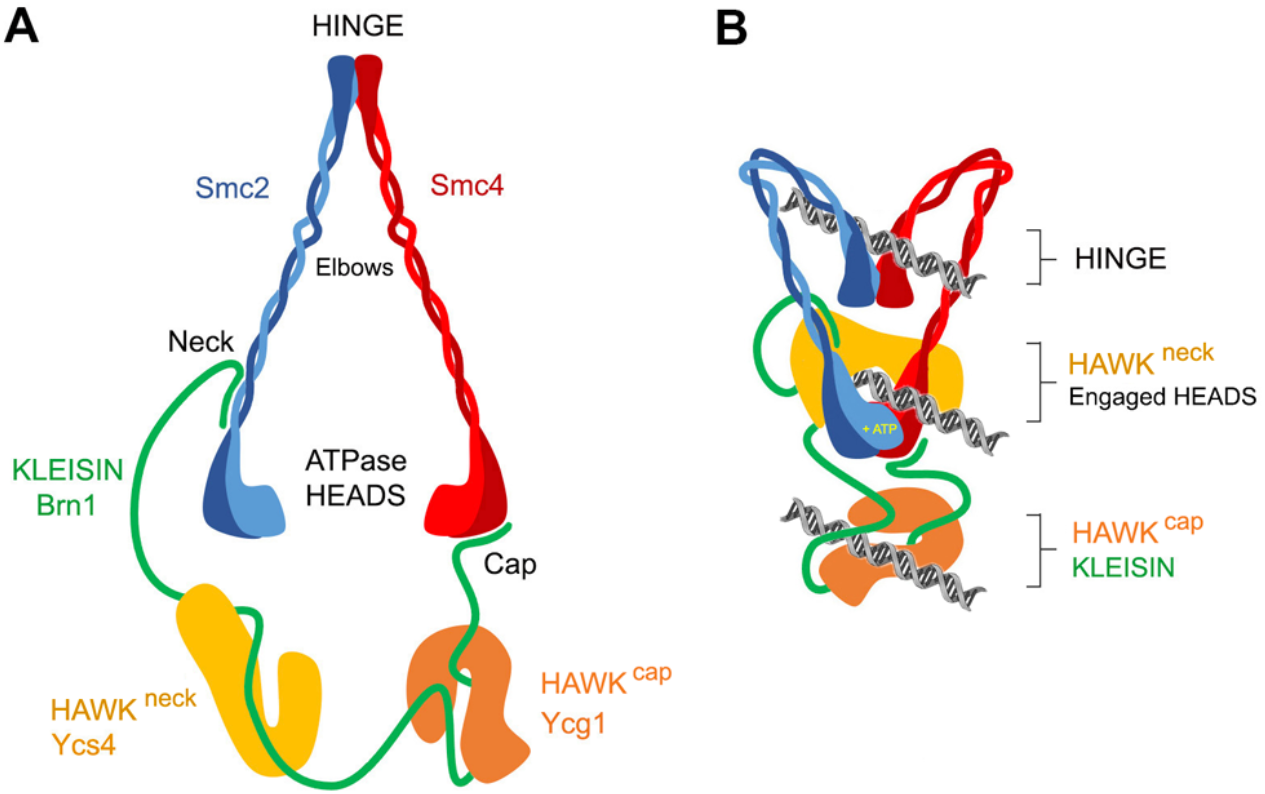
Condensin Architecture. (A) Structure of budding yeast condensin. The Smc subunits (Smc2, Smc4) and the kleisin (Brn1) form a protein ring. Smc2 and Smc4 each fold into a 50 nm long antiparallel coiled-coil that forms a globular “hinge” domain at its apex, whereas the amino and carboxy termini form an ATPase “head” domain at the other extreme. Smc2 and Smc4 dimerize via their hinge domains, while Brn1 forms a long flexible bridge connecting their head domains. The N-terminal domain of Brn1 binds to the coiled-coil “neck” region immediately adjacent to the head of Smc2, and the C-terminal domain to the head tip of Smc4 at the “cap”. The complex is completed by two HAWKs subunits. Ysc4 binds to a Brn1 region proximal to the neck (HAWK^neck^), whereas Ycg1 binds to a Brn1 region proximal to the cap (HAWK^cap^). (B) DNA binding modules and conformational changes. The two ATPase heads engage with each other upon binding a pair of ATP molecules in between them. Interaction of the HAWK^neck^ module with the engaged heads forms a central DNA clamping core. The HAWK^cap^ module binds DNA and encircles it via a kleisin belt. A third DNA binding module is the hinge, which can reach the vicinity of the ATPase heads when the coiled coil arms bend at their elbow.

**Figure S2.**
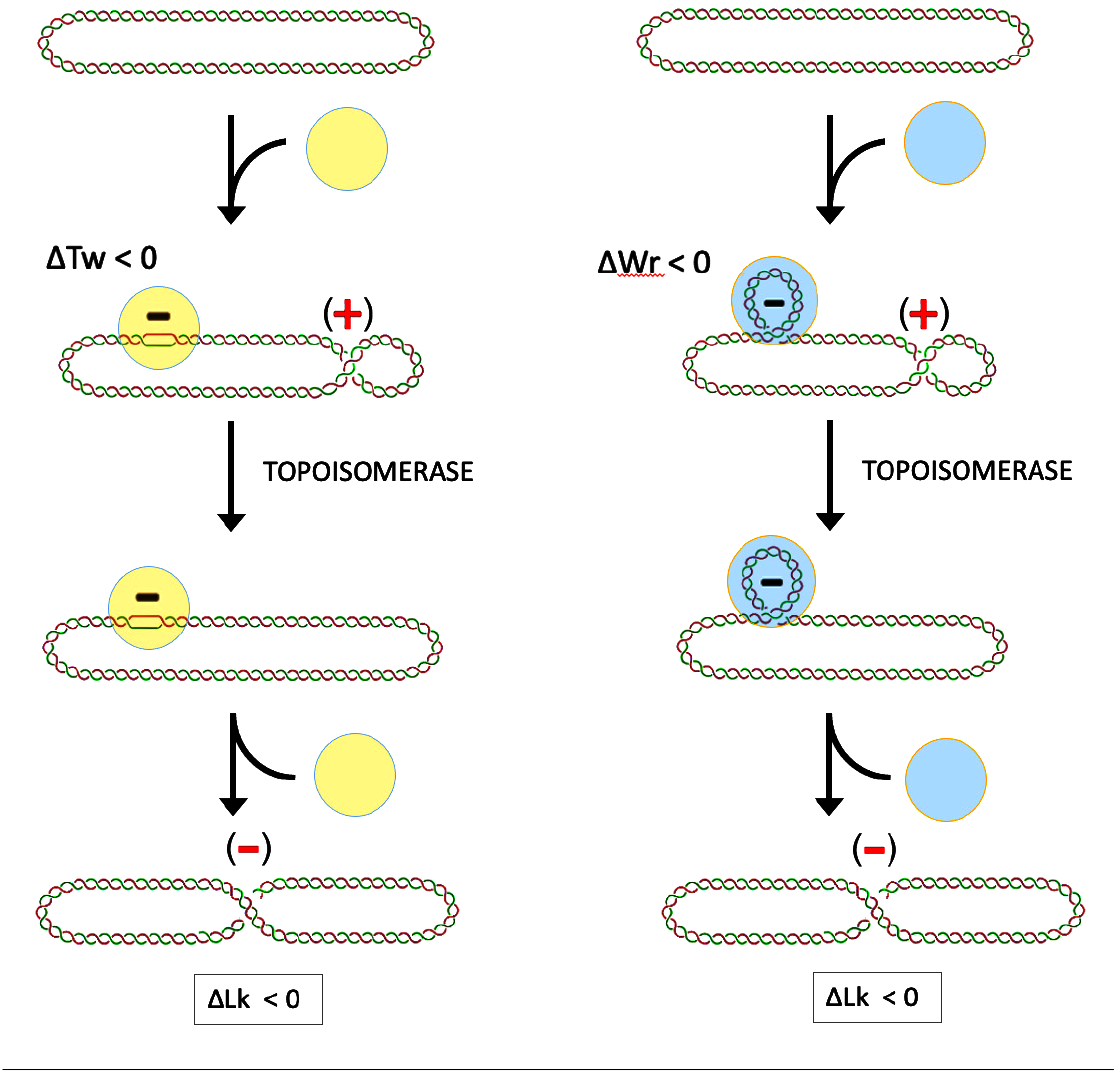
Topological method to assess DNA deformations. The interaction of DNA with any ligand might deform the duplex by producing ΔTw (change of helical winding) and/or ΔWr (non-planar bending). The figure illustrates, on the left, a complex that unwinds the DNA (ΔTw<0) and, on the right, a complex that bends the DNA in a left-handed manner (ΔWr<0). Since the linking number (Lk) of DNA in a closed topological domain equals the sum of DNA twist and writhe (Lk=Tw+Wr), the changes of Tw and/or Wr restrained by the complex will generate compensatory opposite deformations in the form of Tw and Wr. For simplicity, this is illustrated in the figure as an unconstrained (+) supercoil (ΔWr>0). Relaxation of such compensatory (+) supercoil with a topoisomerase will reset the Lk of the DNA. The resulting ΔLk value reflects the ΔTw or ΔWr deformations that were restrained by the bound complex.

**Figure S3.**
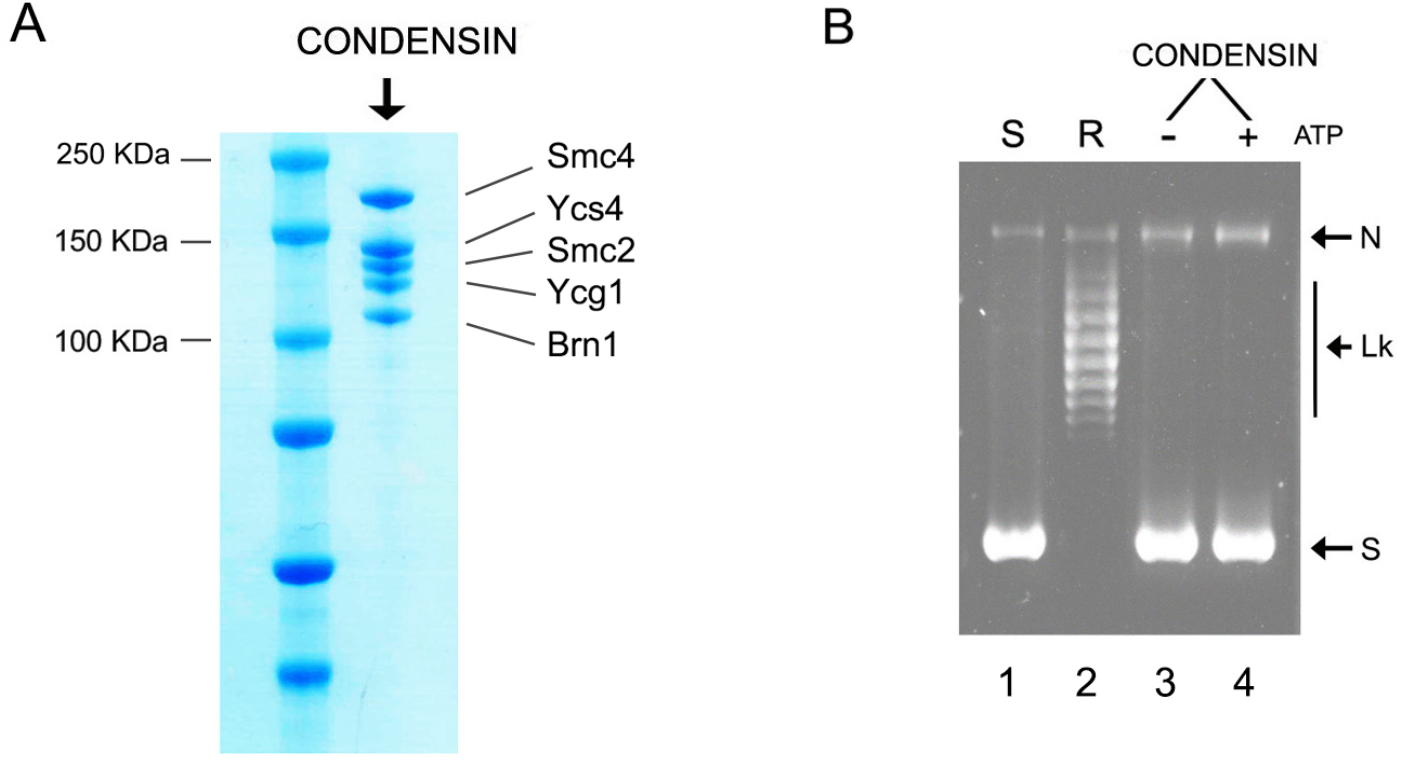
Condensin purification. (A) SDS-PAGE of the purified condensin pentamer (Smc2, Smc4, Brn1, Ycg1 and Ycs4). (B) To verify that the purified condensin complex was free of nuclease and topoisomerase activities, 200 ng of negatively supercoiled DNA plasmid (lane 1) were relaxed with 1 unit of Topo I (lane 2), or incubated for 30 min at 30ºC with 200 ng of Condensin in absence (lane 3) or presence of 1 mM ATP (lane 4). DNA gel electrophoresis was at 2.5 V/cm for 20 h in 0.7% agarose in TBE containing 0.2µg/mL chloroquine. N, nicked DNA; S, negatively supercoiled DNA; Lk, distribution of relaxed topoisomers produced by Topo I.

**Figure S4.**
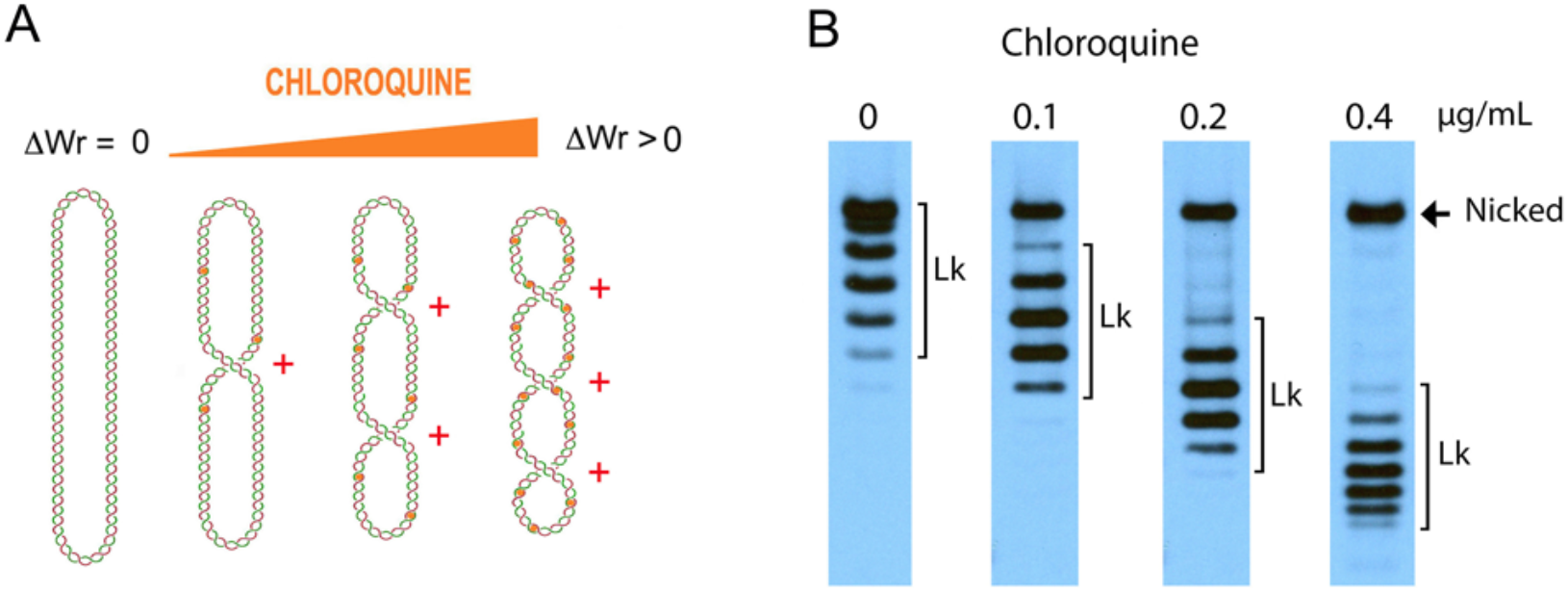
Effect of chloroquine on the gel velocity of Lk topoisomers. (A) Chloroquine intercalation unwinds the DNA double helix (ΔTw<0). In a covalently closed DNA circle, since Lk is constant, such unwinding produces positive helical tension in the chloroquine-free regions and therefore (+) supercoiling (ΔWr>0) of the DNA. Then, increasing the concentration of chloroquine increases the degree of (+) supercoiling. (B) The gel velocity of a covalently closed DNA circle depends of its compaction volume and, therefore, of its degree of supercoiling. Accordingly, the Lk distribution of relaxed DNA will move faster when electrophoresis is conducted in the presence of chloroquine. The figure compares the electrophoretic velocity of the same distribution of Lk topoisomers that run in TBE buffer containing 0, 0.1, 0.2, and 0.4 µg/mL of chloroquine (2.5 V/cm for 18 h in 0.7% agarose).

**Figure S5.**
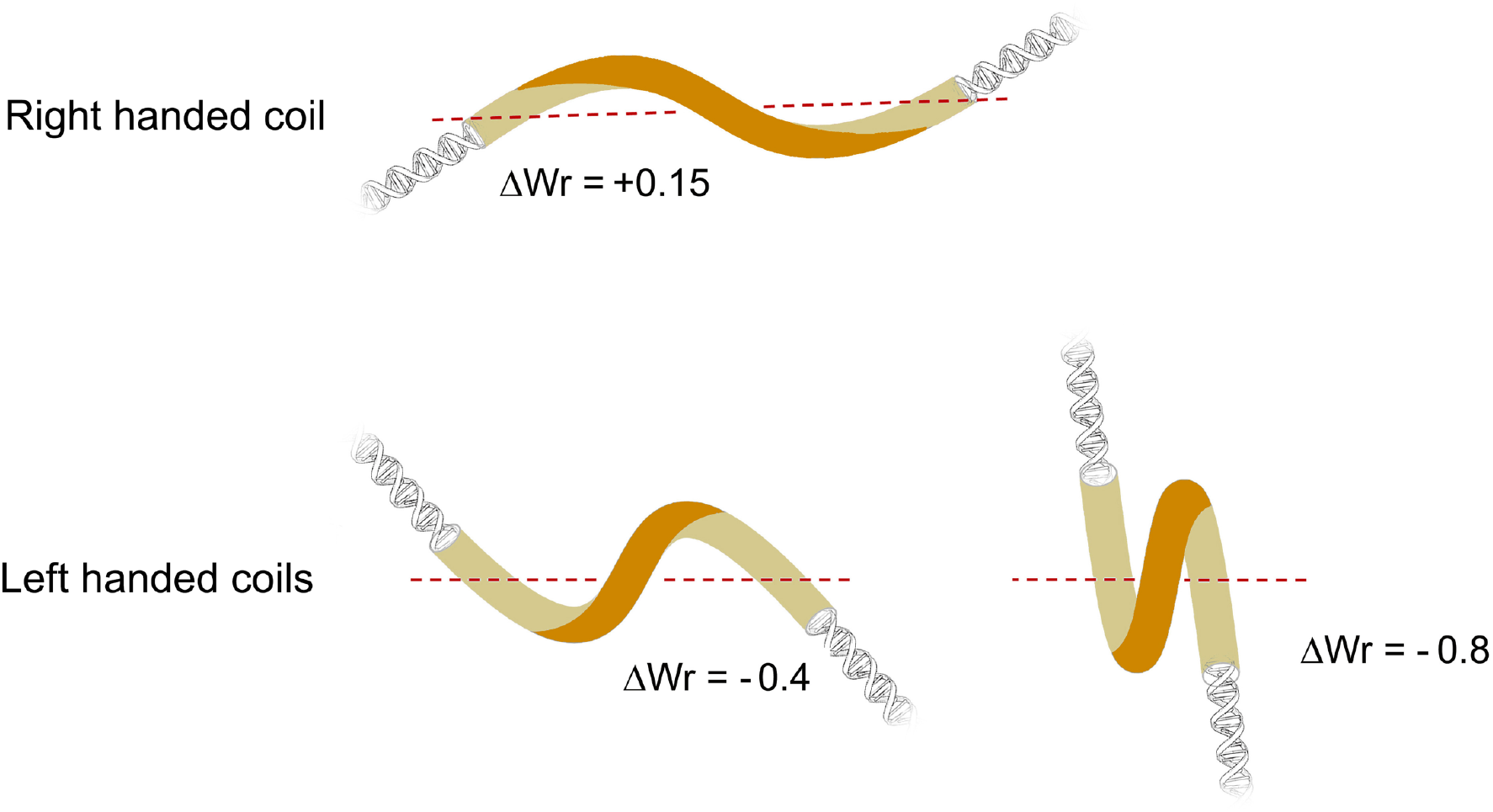
Modeling of DNA deformations as function of ΔWr. Illustrations depict about 120 bp of DNA with ΔWr = +0.15 (right-handed coil) and ΔWr = -0.4 and ΔWr = -0.8 (left-handed coils). Models were done considering that the writhe of a simple coil is given by Wr = 1 – sin ∂, were ∂ is the pitch angle of the coil (Segura et al. 2018).

**Figure S6.**
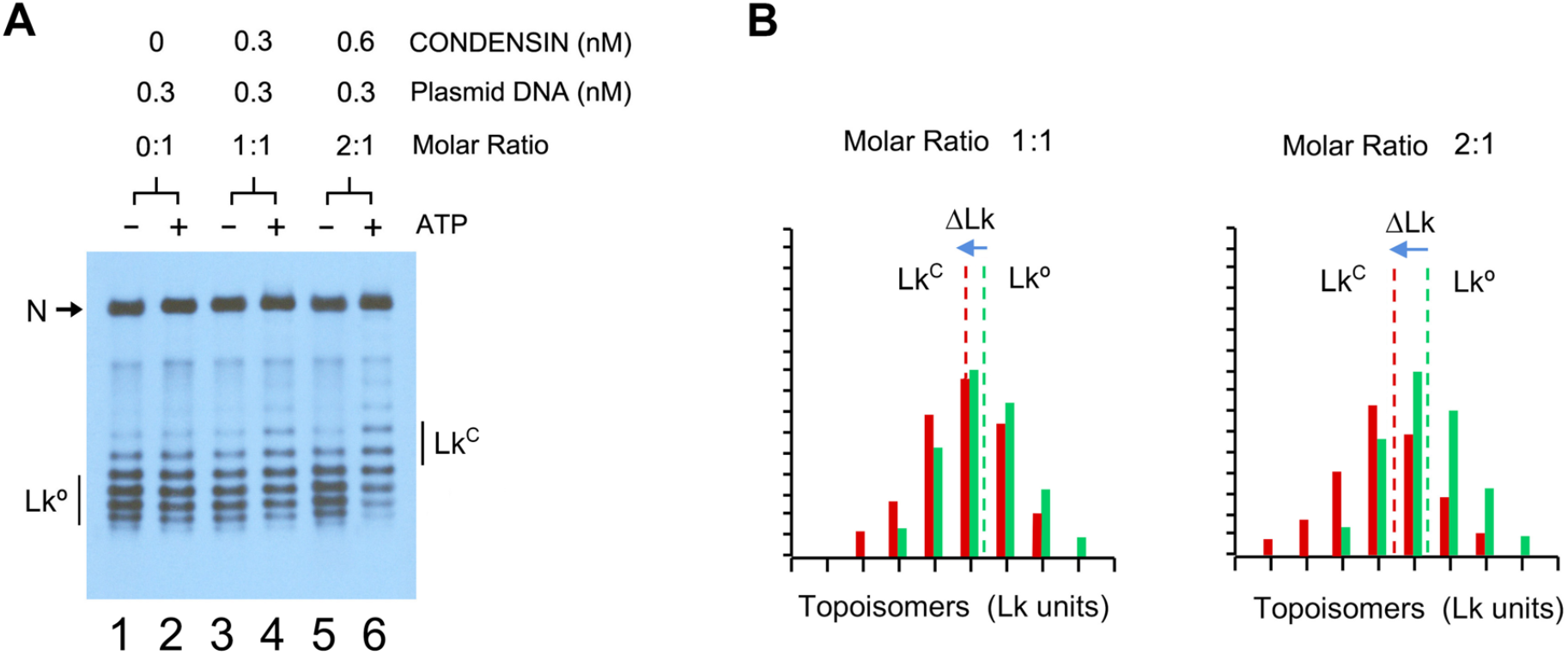
Constraining DNA supercoils with low molar ratios of condensin. (A) Relaxed DNA (0.3 nM), condensin (0, 0.3, 0.6 nM) and Topo I (1 unit) were mixed in 25 mM Tris-HCl pH 7.5, 25 mM NaCl, 5 mM MgCl_2_, 1 mM DTT, with/without ATP (1 mM). Incubations proceeded at 30ºC for 30 min. DNA electrophoresis was at 2.5 V/cm for 20 h in 0.7% agarose and TBE buffer containing 0.4 µg/ml chloroquine. N, nicked circles. Lkº, input distribution of Lk topoisomers of relaxed DNA. Lk^C^, resulting distribution of Lk topoisomers. (B) The histograms compare the relative intensity of individual topoisomers of the Lk distributions in lanes 3 and 4 (condensin:DNA molar ratio 1:1); and lanes 5 and 6 (molar ratio 2:1). Lkº and Lk^C^ denote the midpoint of each Lk distribution, and ΔLk the difference between them.

**Figure S7.**
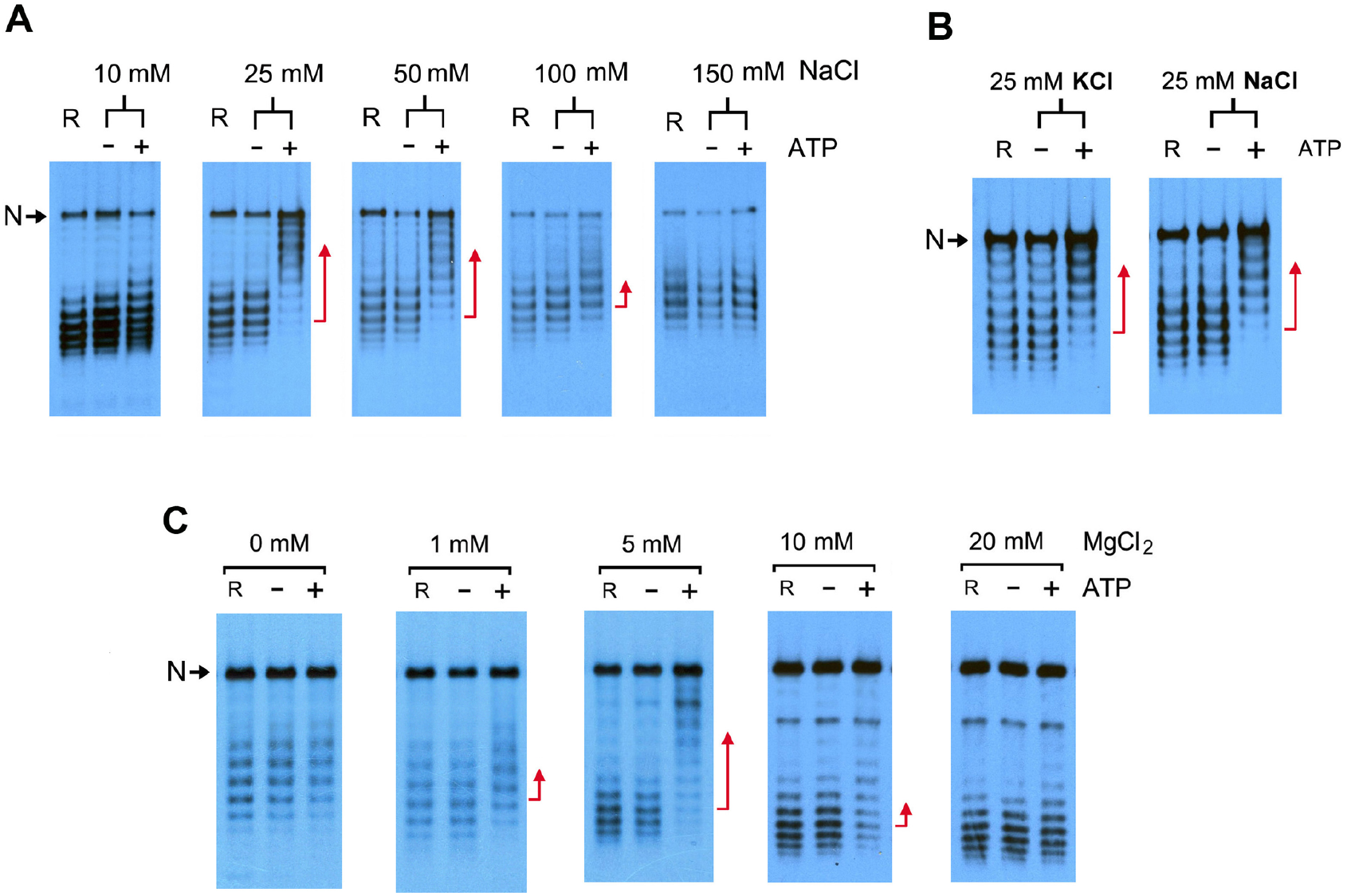
Effect of salt in the condensin capacity to restrain DNA supercoils. (A) Relaxed DNA (0.3 nM), condensin (3 nM) and Topo I (1 unit) were mixed in 25 mM Tris-HCl pH 7.5, 5 mM MgCl_2_, 1 mM DTT, and different concentrations of NaCl (10, 25, 50 100, 150 mM). Mixtures were supplemented without/with ATP (1 mM) and incubated at 30ºC for 30 min. (B) Experiment conducted as in A but containing either 25 mM NaCl or 25 mM KCl. (C) Experiment conducted as in A but containing different concentrations of MgCl_2_ (0, 1, 5, 10, 20 mM). DNA electrophoreses in A, B and C were at 2.5 V/cm for 20 h in 0.7% agarose and TBE buffer containing 0.4 µg/ml chloroquine. Lanes R, DNA relaxed by Topo I (no condensin added). N, nicked circles. Red arrows denote most significant changes of ΔLk restrained by condensin in presence of ATP.

**Figure S8.**
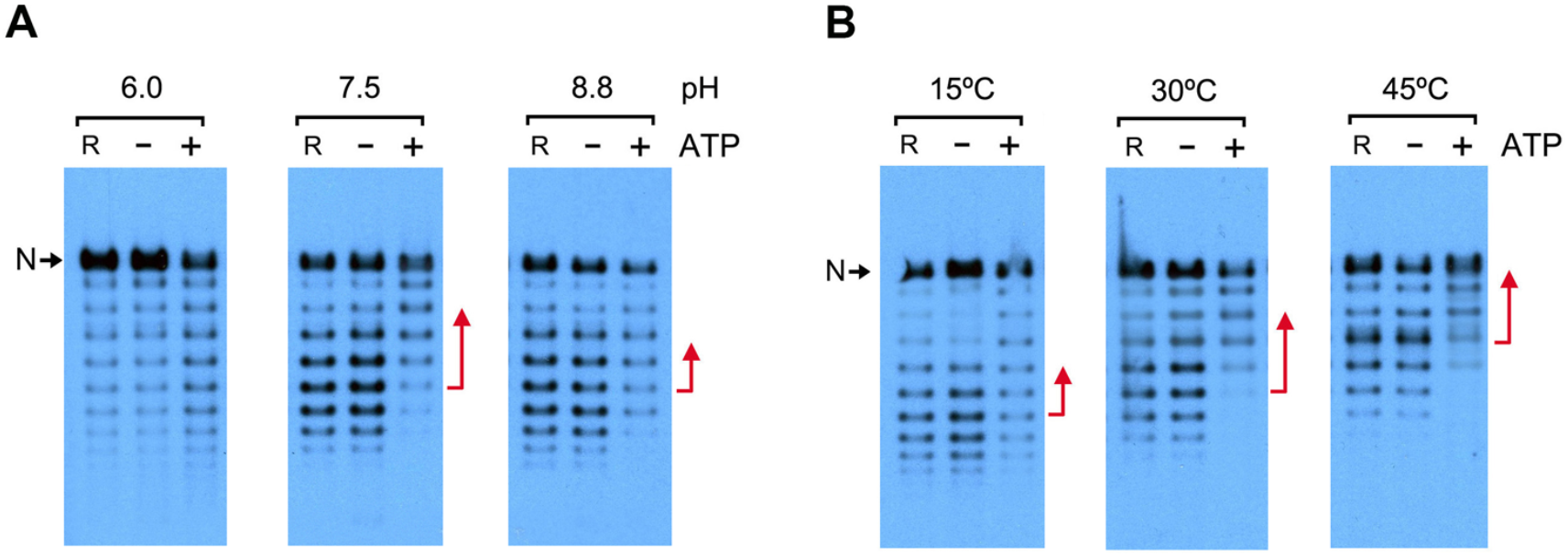
Effect of pH and temperature in condensin capacity to restrain DNA supercoils. (A) Relaxed DNA (0.3 nM), condensin (3 nM) and Topo I (1 unit) were mixed in 25 mM Tris-HCl, 25 mM NaCl, 5 mM MgCl_2_, 1 mM DTT, and pH adjusted to 6.0, 7.5 or 8.8. Incubations supplemented without or with ATP (1 mM) proceeded at 30ºC for 30 min. (B) Relaxed DNA (0.3 nM), condensin (3 nM) and Topo I (1 unit) were mixed in 25 mM Tris-HCl pH 7.5, 25 mM NaCl, 5 mM MgCl_2_, 1 mM DTT. Incubations supplemented without or with ATP (1 mM) proceeded for 30 min either at 15ºC, 30ºC or 45ºC. DNA electrophoreses were at 2.5 V/cm for 20 h in 0.7% agarose and TBE buffer containing 0.2 µg/ml chloroquine. Lanes R, DNA relaxed by Topo I (no condensin added). N, nicked circles. Red arrows denote most significant changes of ΔLk restrained by condensin in presence of ATP. Note that the reference distribution of relaxed Lk topoisomers (Lanes R) also changes by the effect of pH and temperature.

**Figure S9.**
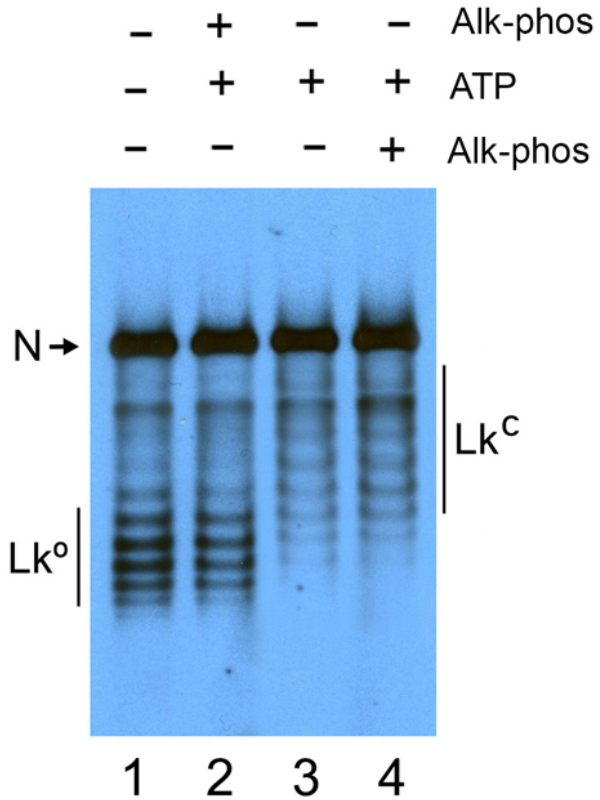
Effect of ATP exhaustion on the condensin capacity to restrain DNA supercoils. Relaxed DNA (0.3 nM), condensin (3 nM) and Topo I (1 unit) were mixed in 25 mM Tris-HCl pH 7.5, 25 mM NaCl, 5 mM MgCl_2_, 1 mM DTT. Incubations proceeded at 30ºC in absence of ATP 1 mM for 10 min (lane 1), in presence of Alkaline Phosphatase and ATP 1 mM for 10 min (lane 2), ATP 1mM for 10 min (lane 3), ATP 1mM for 10 min followed by Alkaline Phosphatase for 60 min (lane 4). DNA electrophoresis was at 2.5 V/cm for 20 h in 0.7% agarose and TBE buffer containing 0.4 µg/ml chloroquine. N, nicked circles. Lkº, distribution of Lk topoisomers of relaxed DNA. Lk^C^, resulting distribution of Lk topoisomers.

**Figure S10.**
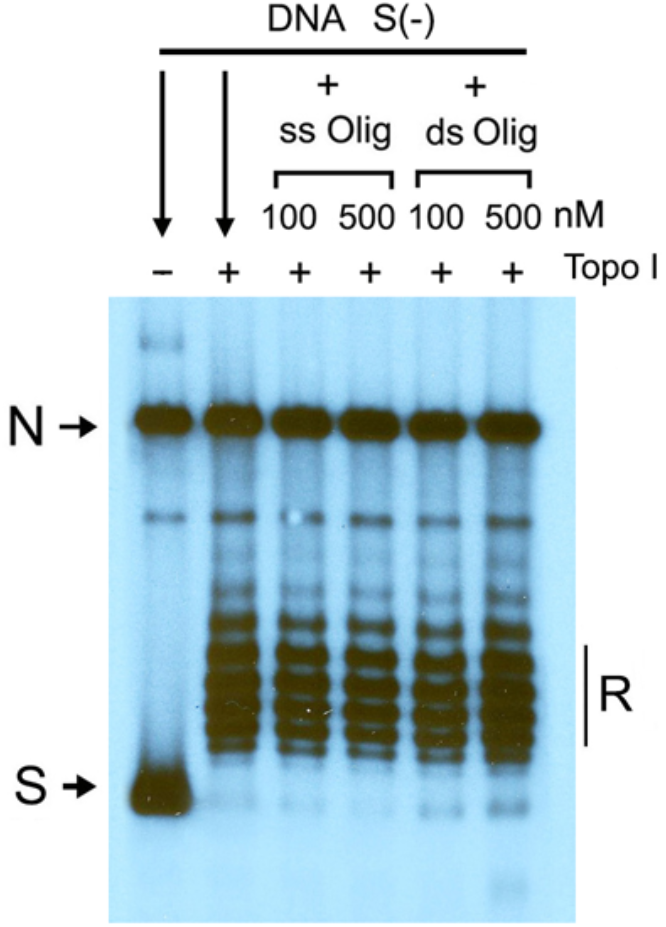
Relaxation of DNA with Topo I in presence of oligonucleotides. Negatively supercoiled DNA (0.3 nM) was mixed with high concentration of ss- or ds-oligos (100 and 500 nM) in 25 mM Tris-HCl pH 7.5, 25 mM NaCl, 5 mM MgCl_2_, 1 mM DTT. Afterwards, Topo 1 (1 unit) was added to the mixtures and incubations proceeded for 10 min at 30ºC. DNA electrophoresis was at 2.5 V/cm for 20 h in 0.7% agarose and TBE buffer containing 0.4 µg/ml chloroquine. S, (-) supercoiled DNA. N, nicked circles. R, relaxed DNA.

**Figure S11.**
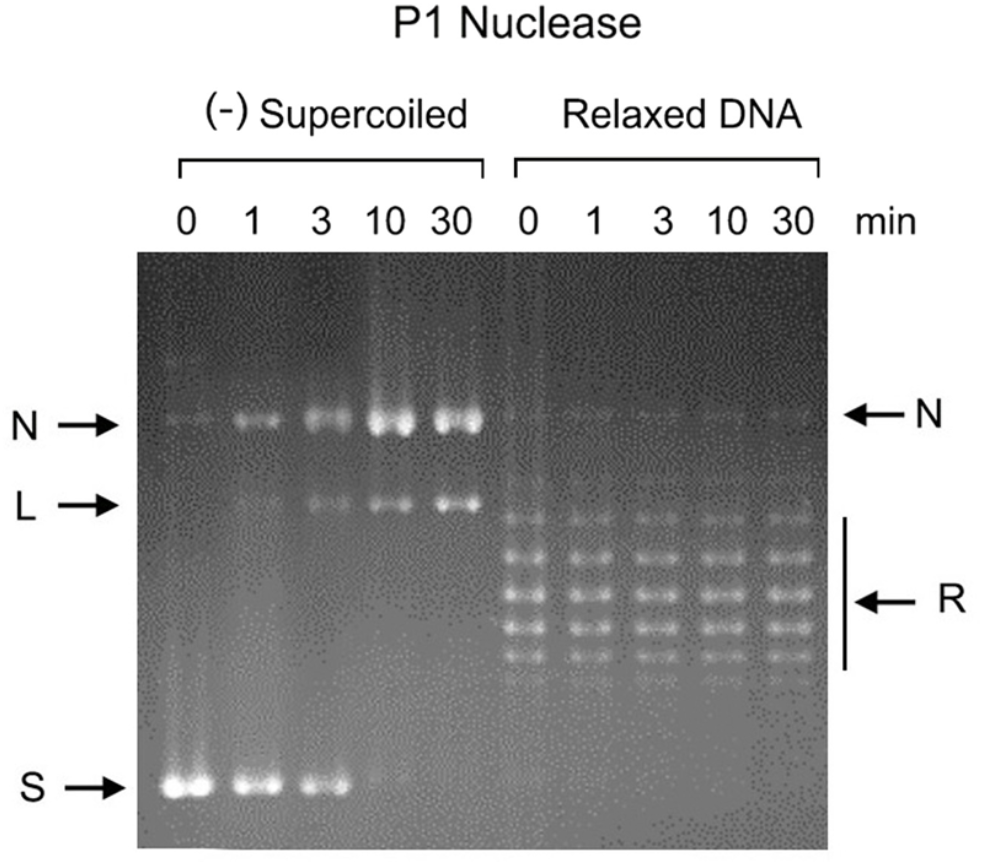
P1 nuclease activity in relaxed and negatively supercoiled DNA. 1µg of either (-) supercoiled or relaxed DNA plasmid were incubated with 10 units of P1 nuclease in 100 µL of 25 mM Tris-HCl pH 7.5, 25 mM NaCl, 5 mM MgCl_2_, 1 mM DTT. Volumes of 20 µL of the ongoing reactions were quenched after 0, 1, 3, 10, 30 min incubation at 30ºC. DNA electrophoresis was at 2.5 V/cm for 20 h in 0.7% agarose and TBE buffer containing 0.2 µg/ml chloroquine. The gel was stained with Ethidium. S, (-) supercoiled DNA. R, relaxed DNA. N, nicked circles. L, linear DNA.

**Figure S12.**
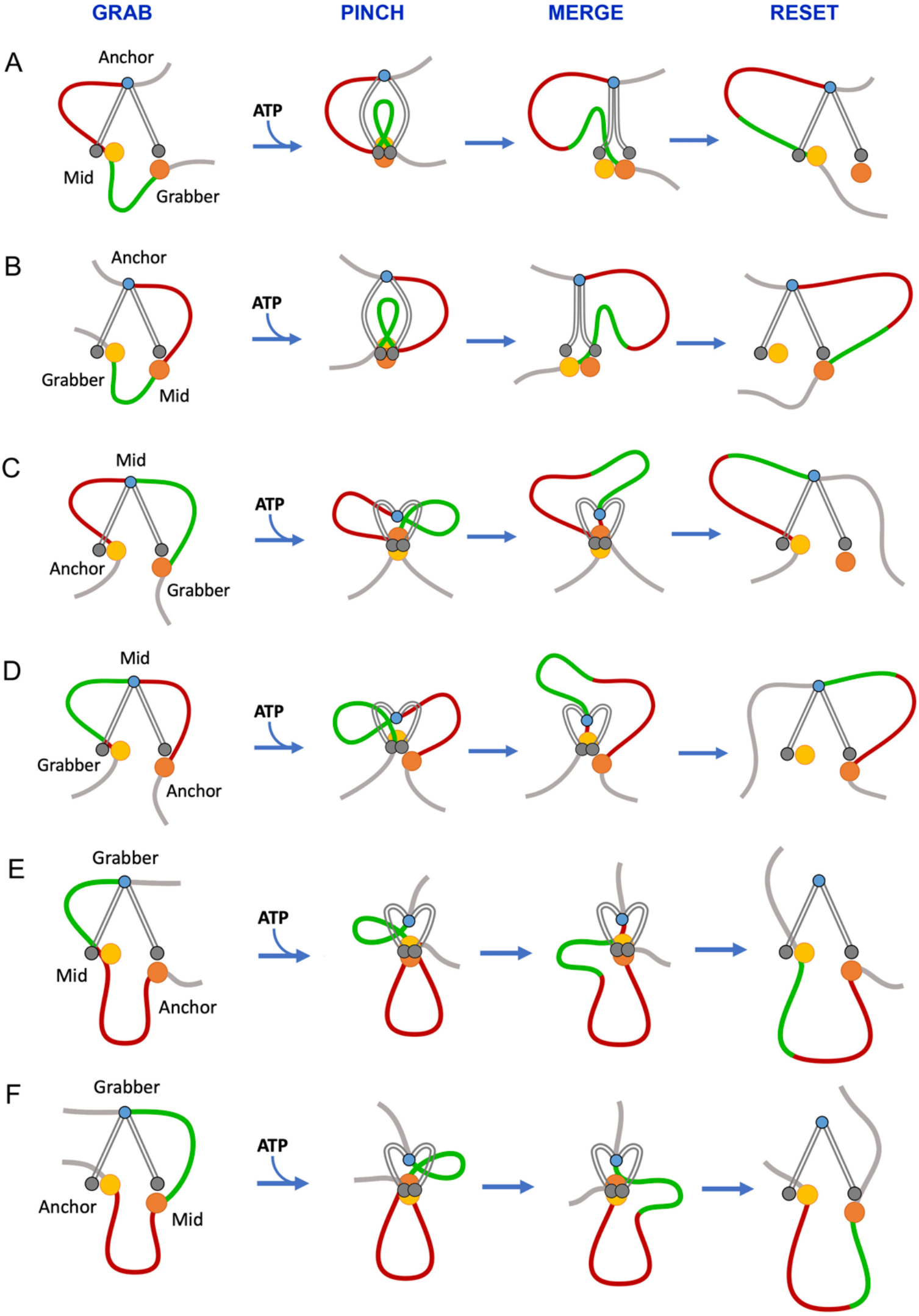
Hypothetical roles of condensin DNA binding modules to perform the “pinch and merge” mechanism. The anchor, mid, and grabber functions are assigned to the three DNA binding modules of condensin in six posible combinations (A-F). For simplicity, the kleisin subunit is not depicted and the HAWKs are represented as spheres (HAWK^cap^ in orange and HAWK^neck^ in yellow). The hinge (in blue) is acting as the anchor in A and B, as the mid site in C and D, and as the grabber in E and F. In all cases, the starting configuration is a loaded complex, in which DNA interacts with the three DNA binding modules. ATP binding produces the engagement of the Smc heads (grey) and the concomitant pinching of a feeding loop (green). ATP hydrolysis triggers the merging process and the reseting of the complex with an enlarged extruded loop (red).

## Notes

### Competing Interest Statement

The authors have declared no competing interest.

